# Fluorescence fluctuation based super resolution microscopy, basic concepts for an easy start

**DOI:** 10.1101/2022.05.06.490863

**Authors:** Alma Alva, Eduardo Brito-Alarcón, Alejandro Linares, Esley Torres-García, Haydeé O. Hernández, Raúl Pinto-Cámara, Damián Martínez, Paul Hernández-Herrera, Rocco D’Antuono, Christopher Wood, Adán Guerrero

## Abstract

Due to the wave nature of light, optical microscopy has a lower-bound lateral resolution limit of about half of the wavelength of the detected light, i.e., within the range of 200 to 300 nm. The Fluorescence Fluctuation based Super Resolution Microscopy (FF-SRM) encompases a collection of image analysis techniques which rely on the statistical processing of temporal variations of fluorescence to reduce the uncertainty about the fluorophore positions within a sample, hence, bringing spatial resolution down to several tens of nm. The FF-SRM is known to be suitable for live-cell imaging due to its compatibility with most fluorescent probes and lower instrumental and experimental requirements, which are mostly camera-based epifluorescence instruments. Each FF-SRM approach has strengths and weaknesses, which depend directly on the underlying statistical principles through which enhanced spatial resolution is achieved. In this review, the basic concepts and principles behind a range of FF-SRM methods published to date are revisited. Their operational parameters are explained and guidance for its selection is provided.

## INTRODUCTION

### BASIC CONCEPTS OF FLUORESCENCE AND THE RESOLUTION LIMIT OF OPTICAL MICROSCOPY

Fluorescence microscopy, in which organic and inorganic fluorophores are employed as molecular dyes, is by far the most popular technique for the observation of biological specimens with molecular specificity (Jacquemet et al., 2020). This level of discrimination is achieved either by labeling cellular components with fluorescent dyes or by linking an engineered fluorescent protein to a molecule of interest (Lichtman and Conchello, 2005).

The Jablonski diagram represents the electronic transitions between energy states during the excitation of a fluorophore (**Figure 1a**). Fluorescence occurs when a fluorophore in a ground electronic state (S_0_) absorbs photonic energy at a specific wavelength range, which promotes an electron to shift to a higher-energy excited state (S_n_). Energy is released upon return of the electron to S_0_, either by non-radiative relaxation or as fluorescence emission in a characteristic spectrum of wavelengths of lower energy. During the excitation cycle, energy is lost, i.e., through molecular kinetics and vibrational relaxation (Vangindertael et al., 2018). The cycle of excitation and fluorescence emission occurs in the range of a few nanoseconds (Lichtman and Conchello, 2005).

**Figure 1.**
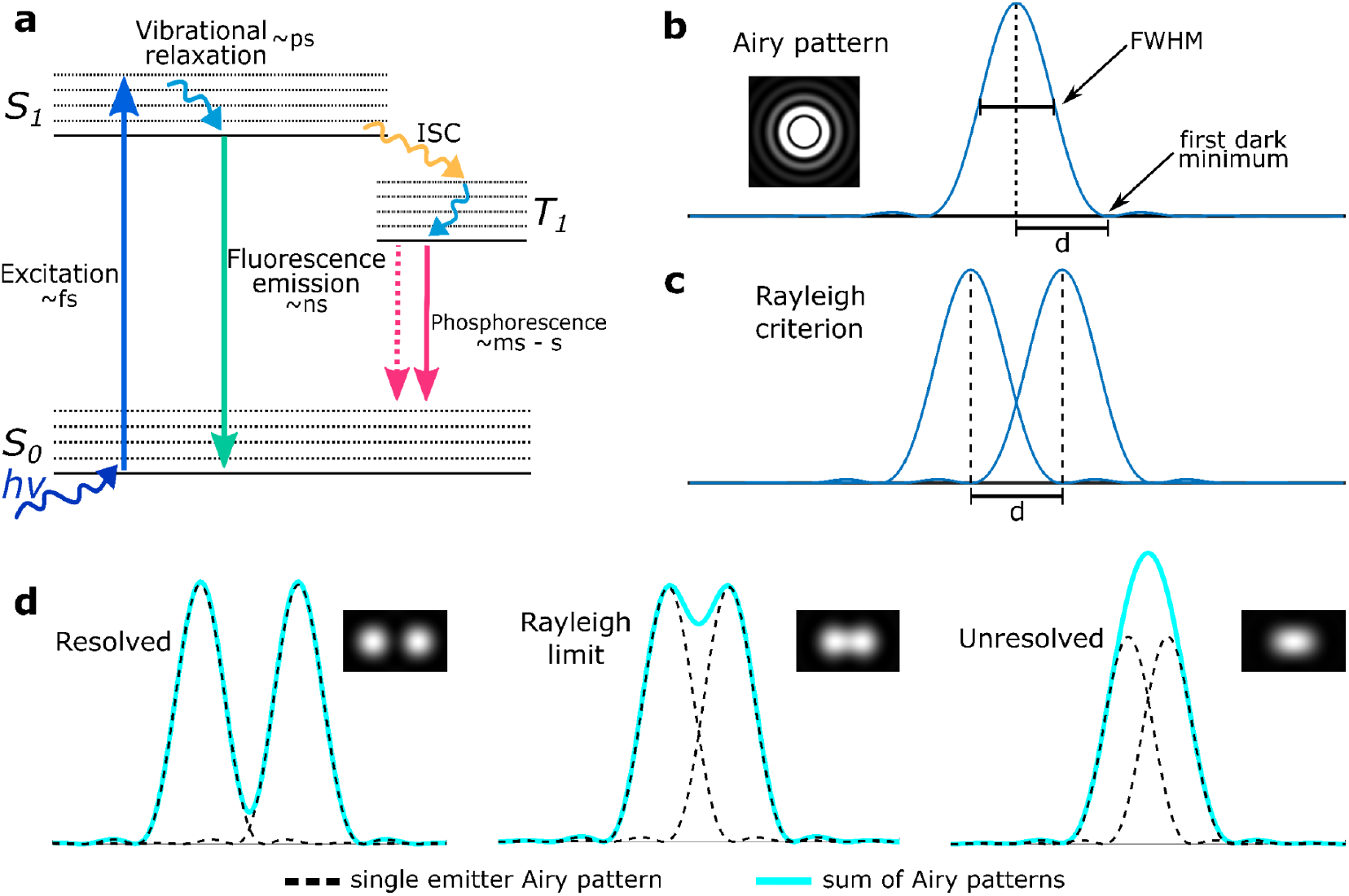
Fluorescence process described by the Jablonski diagram and the spatial resolution limit according to the Rayleigh criterion. **(a)** Simplified Jablonski diagram representing energy states of a fluorescent molecule and their characteristic timescales. The excitation photon bears an energy hν to the fluorophore which causes a transition of an electron from the ground state S_0_ to the excited state S_1_. The excited electrons can return to S_0_ through radiative emission of a photon (fluorescence) or by a non-radiative transition. Less often, the electrons can reach a triplet state (T_1_) by Intersystem Crossing (ISC) on which a change occurs to the spin of the electron. **(b)** Bidimensional representation of the Point Spread Function (PSF) formed by the Airy pattern of one emitter and **(c)** the corresponding Rayleigh criterion for optical resolution when two of these patterns are in close enough proximity to each other. **(d)** Fluorescence distributions of two fluorescence emitters separated at three different distances. From left to right, the emitters are spatially resolved, at the limit of resolution (according to the Rayleigh criterion) or unresolved.

In addition to non-radiative relaxation and radiative fluorescence emission, electrons may cross over into the triplet state (T_1_) by intersystem crossing (ISC), a much longer-lived excited state. From T_1_, an electron may return to the ground electronic state through radiative phosphorescent emission, which typically occurs on the millisecond to seconds time scale and thus, it is easily distinguished from fluorescence emission (Lichtman and Conchello, 2005).

In the wave description of light, the light emitted by a single fluorophore seen through an optical microscope will undergo diffraction as it travels through the microscope optics. Once the light reaches the eye or the microscope detector, it will be observed in the form of a diffraction pattern, a bidimensional representation of the response function of the instrument, also known as the *Point Spread Function* (PSF). The PSF takes the form of a series of concentric disks (the Airy disk pattern) (**Figure 1b**) with a high intensity at its center (Airy disk of order 0) (Vangindertael et al., 2018).

The shape of the PSF depends on the wavelength of the traveling light, the numerical aperture of the microscope and the refractive index of the sample. To compare the resolving power of a microscope, the full width at half-maximum (FWHM) of the PSF intensity profile is calculated (**Figure 1b**); the lower the FWHM is, the greater the resolution of the microscope (Vangindertael et al., 2018; Schermelleh et al., 2019).

According to the Rayleigh criterion, the resolution limit at which two-point emitters cannot be resolved is when the peak of the Airy pattern of one emitter overlaps with the first minimum of the Airy disk of zero-order of the other (R = 0.61 (λem) / NA) (Rayleigh, 1903) (**Figure 1c**). Because the width of the zero-order disk on the PSF equals approximately half of the wavelength of light emitted by the fluorophore, the resolution limit of an optical microscope is approximate ∼200 – 300 nm (**Figure 1d**)

### SUPER-RESOLUTION MICROSCOPY: A NEW ERA OF OPTICAL MICROSCOPY

Fluorescence microscopy enables the observation of biological phenomena with molecular specificity. However, many aspects of these processes remain unknown due to the spatial limit of resolution of the optical microscope. The development of enhanced optical instrumentation, fluorescent probes and mathematical algorithms to overcome this limitation has accelerated considerably in recent years. These efforts have ushered in a new era for the study of nature through light: Optical Super-Resolution Microscopy (SRM).

SRM began with STimulated-Emission-Depletion fluorescence microscopy (STED) (Hell and Wichmann, 1994; Klar et al., 2000) and Structured Illumination Microscopy (SIM) (Gustafsson, 2000). At that time, the maximum spatial resolution achieved with STED was ∼100 nm. Subsequent refinements of these approaches have further pushed the resolution limit to molecular scales (Wegel et al., 2016). However, these techniques require highly-specialized microscopes, whose cost and complexity have limited their application to a reduced portion of the bioimaging community (Jacquemet et al., 2020).

With the development and applications of suitable photobleaching-resistant and photoconvertible fluorophores, a set of SRM techniques based on the localization of blinking fluorophores and without the requirement of a STED microscope setup, Single Molecule Localization Microscopy (SMLM), appeared (Jacquemet et al., 2020). Nevertheless, some SRM variants based on STED, such as REversible Saturable Optical FLuorescence Transitions (RESOFLT) and MINFLUX require fluorophores with a drastic fluorescence fluctuation such as blinking (Hofmann et al., 2005; Balzarotti et al., 2017).

SRM based on SMLM can construct a super-resolved image because only a few emitters (around 10 emitters μm^-2^) are collected on a single frame, diminishing the probability that the fluorescence distribution of any two emitting fluorophores overlap (Lelek et al., 2021). The position of those single emitters can be estimated with greater precision through the fitting of a Gaussian function to identify the centroid position. Since only a small fraction of the total fluorophores are emitting, it is imperative to acquire several thousands of frames of the observed field. Depending on the exact characteristics of the SMLM protocol employed, the maximum resolution achievable has been reported to be in the range of ∼ 5 nm (Schnitzbauer et al., 2017; Lelek et al., 2021).

SMLM requires the use of fluorophores capable of transit between prolonged emitting ‘on’ and non-emitting ‘off’ states while avoiding the irreversible photobleached state (**Figure 2a**). With adequate sample preparation (e.g. fluorophore selection, oxygen scavenging buffers, and laser power selection) the probability of the fluorophore to transit between the ‘on’ and the ‘off’ state can be optimized (Minoshima and Kikuchi, 2017; Lelek et al., 2021). Such tuning is highly experiment-dependent, and much time must be invested to optimize the acquisition protocol for each set of experimental conditions and microscope employed. Disadvantages of SMLM techniques include the need for high power laser illumination (from ∼62 to 7.8 kW/cm^2^) (Lin et al., 2015), sample drift in x, y and z position, detectable fluorophores at the last set of acquired images, photodamage (Jacquemet et al., 2020).

**Figure 2.**
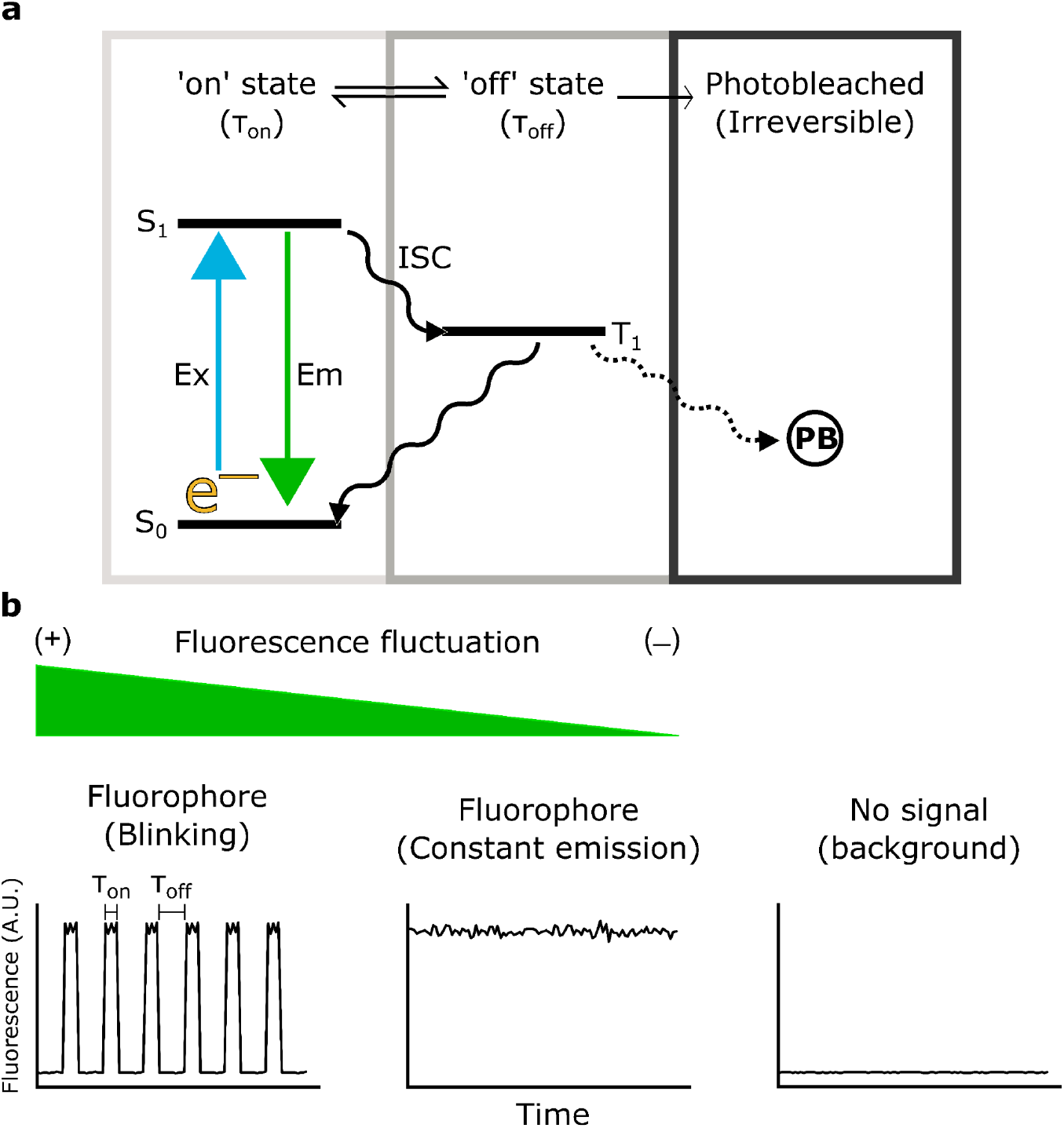
Fluorescence intermittency. **(a)** Naturally, the fluorophores change between the ‘on’ and the ‘off” state, which arises a formal blinking pattern. In some cases, a set of fluorophores transition from the triplet state to a photobleached irreversible state. Under the same fluorophore dynamics, a constant fluorescent emission is characterized by a short ‘off’ state but it also has fluorescence fluctuations as a minor contribution. **(b)** Fluorescence fluctuations contributions depend on fluorophore emission state. Formal blinking fluorophores will contribute more fluctuations in comparison to constant emitting fluorophores, every camera will detect low levels of fluorescence even with no fluorophores present.

On the other hand, Fluorescence Fluctuation based Super-Resolution Microscopy (FF-SRM) relies on the statistical analysis of fluorescence fluctuations over time (Dertinger et al., 2009; Cox et al., 2011; Yahiatene et al., 2015; Agarwal and Macháň, 2016; Gustafsson et al., 2016; Li et al., 2018; García et al., 2021). It can be used for blinking fluorophores in which the fluctuations are remarkable or with constant emitting fluorophores where the fluctuations are generated by intrinsic variations of the fluorescence and not by prolonged ‘off’ states. In both cases, the fluorescence detected must be above the background signal (Moeyaert, Vandenberg and Dedecker, 2020) (**Figure 2b**). The main difference between the SMLM and FF-SRM methods is that the first algorithms are based on the localization of the emitter making the SMLM more precise at the cost of more complex sample preparation and image acquisition (Geissbuehler, Dellagiacoma and Lasser, 2011).

Most of the FF-SRM methods are compatible with epifluorescence, confocal and Total Internal Reflection Fluorescence (TIRF) microscopy, as long as image sampling satisfies the Nyquist-Shannon criteria, meaning that the effective pixel size should be at least half that of the PSF (Vangindertael et al., 2018). As shown in **Figure 3a**, FF-SRM methods have been constantly improved, generating a family of FF-SRM tools that can achieve a spatial resolution at the nanoscopic scales.

**Figure 3.**
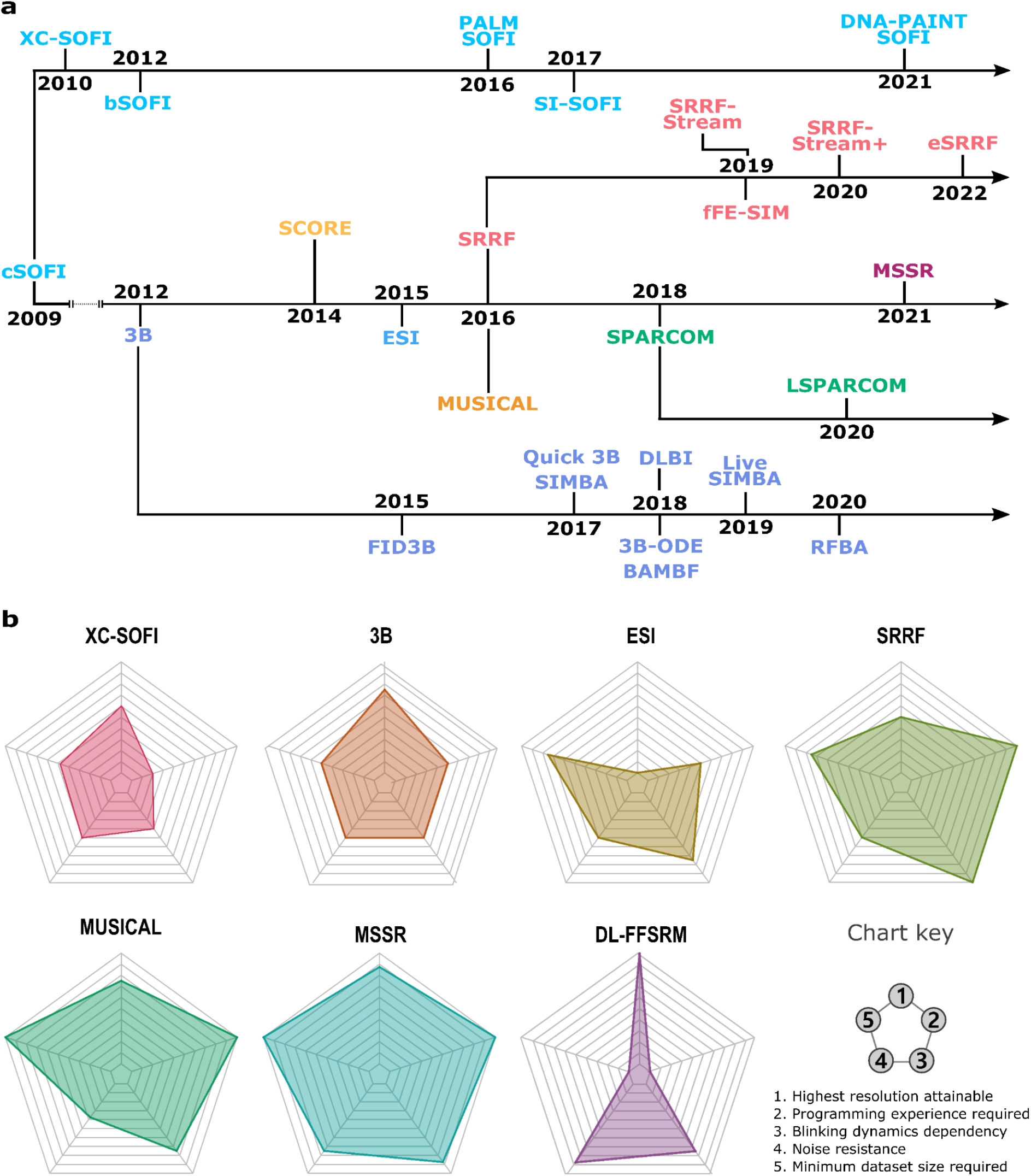
History and objective parameters related to fluorescence fluctuation based super resolution microscopy (FF-SRM) **(a) Timeline of FF-SRM methods release**. The methods located on the central branch represent their first publication and implementation. A different color is assigned to each method and each branch from the original method contains the corresponding improved versions following the seeding color scheme. **(b) Comparative web-diagrams showing the strengths and weaknesses of the discussed FF-SRM methods**. The most relevant experimental aspects to consider before FF-SRM method selection are displayed in the chart key panel. Abbreviations: XC-SOFI: Cross-cumulants Stochastic Optical Fluctuation Imaging: 3B: Bayesian analysis of Blinking and Bleaching; ESI: Entropy-based Super-resolution Imaging; SRRF: Super Resolution Radial Fluctuation; Multiple Signal Classification Algorithm; MSSR: Mean-Shift Super Resolution.

All FF-SRM methods work with relatively similar fundamentals; they gather nanoscopic information through an analysis of temporal fluorescence intermittency (Dertinger et al., 2009; Cox et al., 2011; Yahiatene et al., 2015; Agarwal and Macháň, 2016; Gustafsson et al., 2016; Li et al., 2018; García et al., 2021). Each one has particular requirements for its image acquisition strategy, which include aspects of sample preparation and imaging parameters employed such as the minimum number of frames, optical and software parameters, image amplification, among others (Jacquemet et al., 2020). **Figure 3b** summarizes the strengths and weaknesses of the FF-SRM methods described in this review.

#### **S**tochastic **O**ptical **F**luctuation **I**maging

Published in 2009, Stochastic Optical Fluctuation Imaging (SOFI) was the first FF-SRM method (Dertinger et al., 2009) and is considered the founder of this family. SOFI is grounded on the analysis of the temporal dynamics of fluorescence fluctuations at nanoscopic scales. In this section, we will cover the first and second SOFI implementations, published by (Dertinger et al., 2009) and (Dertinger et al., 2010), respectively. To differentiate one work from another, we will refer to them as classic SOFI (cSOFI) and cross-cumulants SOFI (XC-SOFI), respectively.

cSOFI embraces the analysis of a temporal stack of fluorescent images. It considers a fluorescence image as a digitized collection of *N* emitters, located at **r**_k_ positions (with k=1, 2, …, N), with constant molecular brightness. At each pixel, the fluorescence signal fluctuates in a time-dependent manner, due to the fluorescence intermittency of the emitters it harbors. Within a single fluorescence image, each emitter is considered to be convolved with the PSF of the optical system. Hence, a cSOFI experiment encompasses the analysis of a temporal sequence of images gathered from a static scene, in which the fluorescence fluctuation contains nanoscopic information not available in the spatial domain of a single diffraction-limited image (**Figure 4a**).

**Figure 4.**
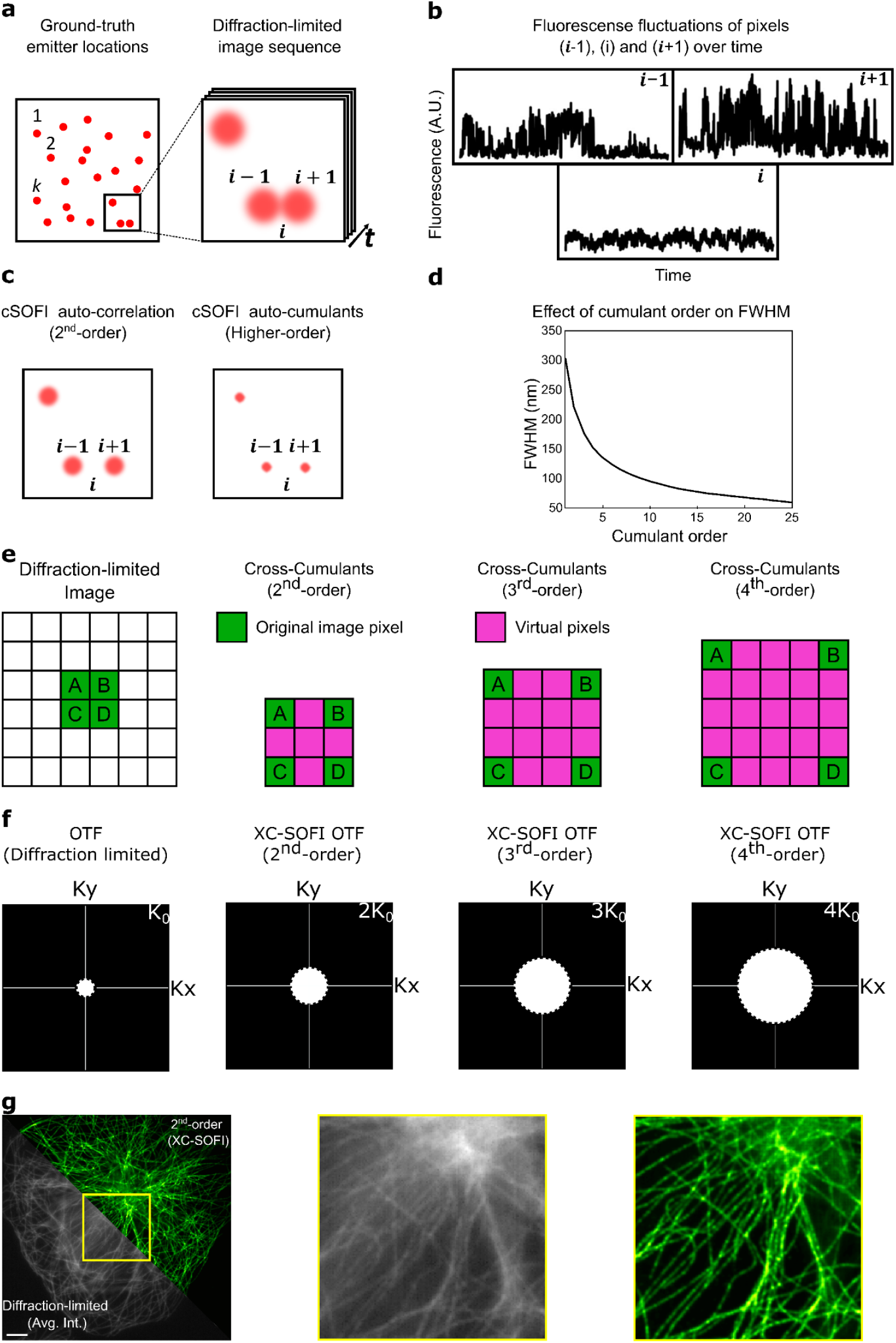
cSOFI and XC-SOFI. (**a**) Original single emitter positions (Ground-truth) and its respective diffraction-limited image sequence. (**b**) Fluorescence fluctuation signal registered across time corresponding to pixels (*i*-1), (*i*) and (*i*+1). (**c**) Super-resolved cSOFI of second and higher SOFI orders. (**d**) Resolution limit of cSOFI. Based on (Dertinger et al., 2009) (**e**) Generation of novel pixels by the cumulants in XC-SOFI. (**f**) PSF in Fourier Space (OTF) diffraction limited and extended by several SOFI orders. (**g**) Super-resolved microtubules by 2^nd^-order XC-SOFI. Image was reconstructed using XC-SOFI within Localizer, Igor Pro 8.0.4 (Dedecker, Duwé, et al., 2012). Data obtained from (Sage et al., 2019). Scale bar: 5 µm.

An ideal sample for a cSOFI experiment is one on which the fluorophores are spatio-temporally static (*i.e*. no sample drift) and fluorescence fluctuations due to transition through excited states are the main cause of change in the fluorescence signal. The fluctuations from the background (pixel *i*) are different from the fluorescence fluctuations, hence, they can be separated (**Figure 4b**). To separate the fluorescence from the background, cSOFI seeks for temporal self-similarities of the signal of each pixel with itself at a time τ, this is achieved through computing the temporal autocorrelation function or related mathematical treatments, as described in (Dertinger et al., 2009). This is the elemental form of cSOFI, which improves the resolution by 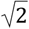 and it is called second-order SOFI (**Figure 4c**).

For further resolution enhancement, it is possible to compute cSOFI of higher orders (*n* ≥ 3). This approach does not auto-correlate the fluorescence fluctuation signal, instead, it computes further statistical descriptors of the fluorescence dynamics called temporal cumulants, which are similar to the statistical moments (Mendel, 1991). Higher-order auto-cumulant cSOFI improves the resolution by a factor of 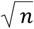, where *n* is the order of cSOFI (**Figure 4c**). In theory, a 4^th^-order cSOFI generates a 2-fold improvement in resolution 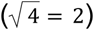 and a 16^th^-order cSOFI can achieve a 4-fold resolution increase 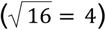. Despite the order of cSOFI used, it has been demonstrated that the experimental cSOFI resolution is ∼ 60 nm (**Figure 4d**) (Dertinger et al., 2009).

The increase in resolution in cSOFI can be scored by measuring the reduction of the PSF as a function of the order of sSOFI. However, a limitation of cSOFI is that the image pixel size is constant between the input diffraction-limited dataset and the super-resolved image. This issue imposes a boundary to the achievable resolution (Dertinger et al., 2010). If the pixel size of the diffraction limited dataset is oversampled (more than two pixels covering the full width of the PSF), higher orders of cSOFI will deliver images with enhanced spatial resolution. However, if the pixel size of the diffraction limited dataset is at the Nyquist-Shannon sampling criterion, or bigger than it (less than two pixels covering the full width of the PSF), the gain of resolution provided by higher orders of cSOFI will be obscured, or even lost, due to the undersampling of higher spatial frequencies.

To generate a super-resolved image with more pixels, XC-SOFI discards the auto-cumulants analysis and employs cross-cumulants instead (Dertinger et al., 2010). This mathematical model approach generates novel virtual pixels that increase with the order of XC-SOFI (**Figure 4e**). The amplification of the original image generates a checkerboard effect. To circumvent this problem, the fluorescence intensity assigned to each virtual and original pixel is modulated by the distance factor. At this point, a preliminary image of XC-SOFI has already been generated. The XC-SOFI algorithm finalizes with a deconvolution step that balances the contrast of the images, facilitating the observation of the gained spatial frequencies.

In the Fourier space, the objective lens of a microscope acts at a finite aperture which limits the spatial frequencies of the diffraction limited image. This can be represented by the Optical Transfer Function (OTF), which is itself the Fourier transform of the PSF. The Abbe criterion and the numerical aperture of the lens (NA) can be directed linked by the maximum observable spatial frequency K_0_ = 2 NA / λ_em_, which can be conveniently represented by a circle of radius K_0_ within the real part of the OTF (also called the Modulation Transfer Function) (**Figure 4f**, white dashed circles). Reviewed in (Dertinger et al., 2010; Geissbuehler, Dellagiacoma and Lasser, 2011; Vangindertael et al., 2018; Pawlowska et al., 2022).

The deconvolution step used by XC-SOFI is a Wiener deconvolution (Sibarita, 2005; Dertinger et al., 2010). **Figure 4f** shows the maximal spatial frequency obtained after each order of XC-SOFI followed by Wiener deconvolution. This strategy extends the K_0_ radius by the SOFI order (K_0_ · *n*). The resolution enhancement of XC-SOFI is *n* rather than 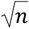, as in cSOFI (Dertinger et al., 2010; Geissbuehler, Dellagiacoma and Lasser, 2011; Pawlowska et al., 2022).

XC-SOFI was used for live imaging of HeLa cells along with a reversible photoswitching protein called Dronpa, compared to previous SOFI implementations that used indirect fluorescent tagging and a formal blinking fluorophore (Dedecker, Mo, et al., 2012).

cSOFI and XC-SOFI have shown capabilities for 3D SRM by analyzing Z-stack data sets acquired on a widefield microscope (Dertinger et al., 2012). Moreover, XC-SOFI has also been used for dual-color SRM (Gallina et al., 2013). In 2014, bSOFI was used in images acquired by a custom microscope capable of multi-channel and simultaneous multi-plane imaging of fixed 3D mitochondria networks in C2C12 cells, and live-cell imaging of HeLa cells (Geissbuehler et al., 2014).

A combination of PhotoActivation Localization Microscopy (PALM) (Betzig et al., 2006) with bSOFI (Geissbuehler et al., 2012) was reported for the imaging of focal adhesion dynamics, and by combining both methods on the same data, the authors generated an improved SR image in comparison with each super-resolution methods applied independently (Deschout et al., 2016). Meanwhile, another merge of techniques, now at the fluorophore level, between bimolecular fluorescence complementation (BiFC) (Kodama and Hu, 2012) with Dronpa resulted in a novel strategy for determining protein-protein interactions in live cells, with the application of XC-SOFI analysis to obtain the nanoscopic position of the interacting proteins (Hertel et al., 2016).

SOFI has been used in several biological models and the theory has been developed far beyond the seminal publication of cSOFI. In 2012, Geissbuehler et al. (2012) restructured the math of XC-SOFI itself, as in balanced SOFI, which in addition to the super-resolved image, can generate quantitative molecular information, such as molecular brightness, on-time ratio, and density of fluorophores.

SOFI does not demand controllably blinking fluorophores, due to its capabilities to discern between lower fluorescence fluctuations and background fluctuations over time. Nonetheless, the greater the difference between the background and a more blinking fluorescence signal, the better the super-resolved image will be (Geissbuehler, Dellagiacoma and Lasser, 2011; Moeyaert, Vandenberg and Dedecker, 2020).

The number of frames required to reconstruct a super-resolved scene using SOFI depends on the SOFI variant, and on the SNR and the desired SOFI order to calculate, i.e., 3000 to 5000 frames for 3^rd^-order and >10,000 frames for 4^th^-order. Like other SRM methods, SOFI is not artifact-free and there should be caution when the super-resolved image is generated. Based on the information presented here, for the microscopists interested in SOFI, we encourage the reading of an exhaustive SOFI review (Pawlowska et al., 2022).

#### **B**ayesian analysis of **B**linking and **B**leaching

The Bayesian analysis of Blinking and Bleaching (3B) estimates the position of the molecules using Bayesian inference, factorial Hidden Markov Chains, and Markov Chain Monte Carlo (MCMC) sampling (**Figure 3a**) (Cox et al., 2011). 3B works with a high density of fluorophores, i.e. each frame can contain overlapping fluorophores, and may require as few as 300 images to compute an SRM image (Cox et al., 2011; Xu et al., 2015). For this reason, 3B can be used for both fixed or live cell imaging on wide-field images of samples expressing typical levels of fluorescent proteins, achieving a spatial resolution of ∼ 50 nm.

In the 3B analysis, each fluorophore is modeled as a Markov chain transiting between three possible states: S_0_, S_1_, and S_2_ (**Figure 5b**). S_0_ and S_1_ are excited states. Emission of fluorescence takes place at the S_0_ state, whereas the S_1_ state encompases transition towards non-fluorescent excited states such as the triplet state. The S_2_ state indicates that the fluorophore has reached an irreversible dark state (photobleached). The transition between states is characterized by transition probabilities (P_n_ where n = 1, …, 6 as depicted in **Figure 5b**). Throughout the image acquisition, the fluorophores in S_0_ or S_1_ can persist in the same state (P_1_ and P_4_, respectively), or switch between S_0_ to S_1_ (P_2_ and P_3_). Fluorophores that have entered the S_2_ (P_5_), cannot regress to S_0_ or S_1_ (P_6_). These transitions define the transition probability matrix which is related to the temporal stochasticity of fluorescence dynamics.

**Figure 5.**
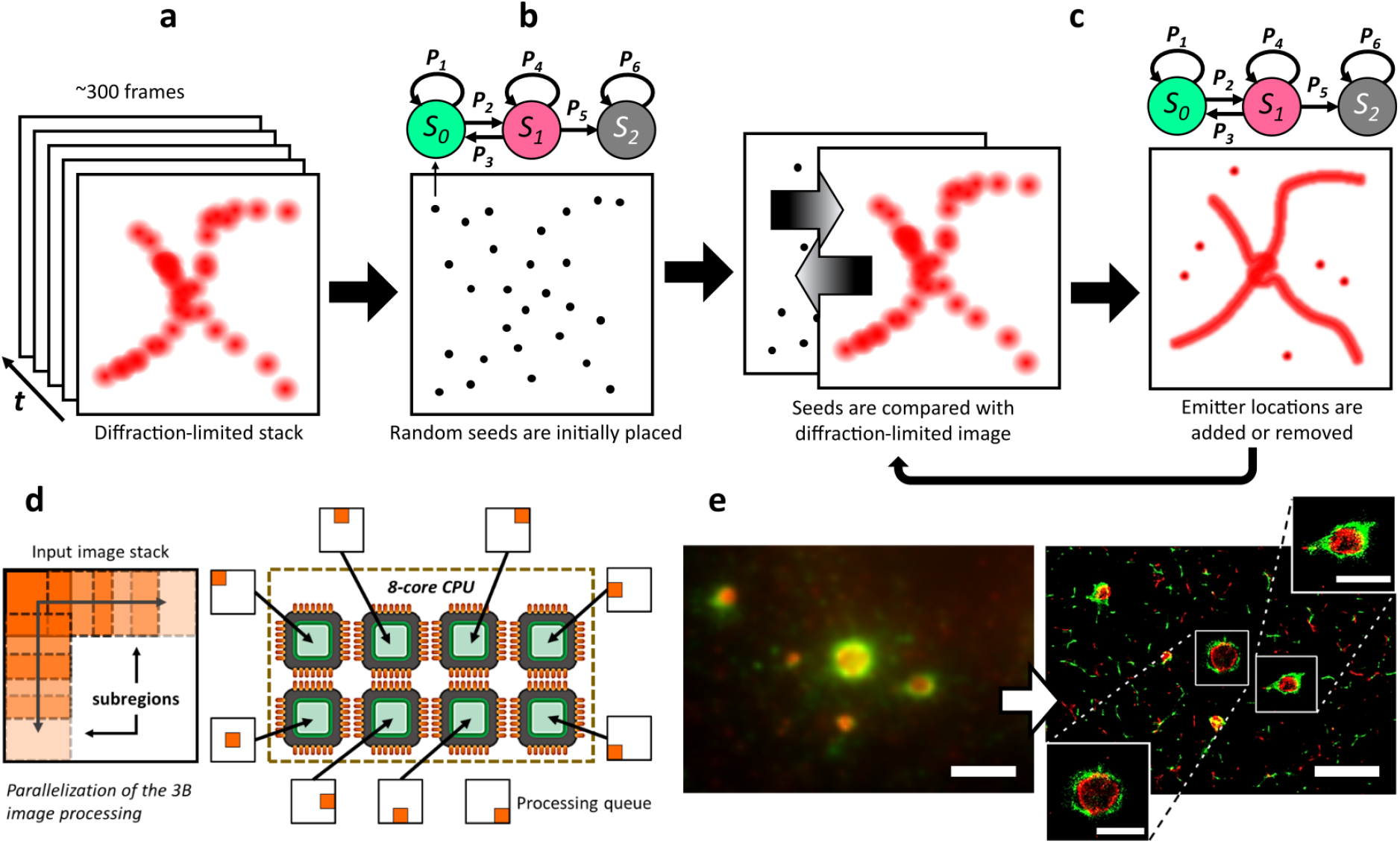
Bayesian analysis of Blinking and Bleaching (3B) microscopy. **(a)** The input data is a diffraction-limited stack of fluorescence images containing ensemble fluorophores undergoing a time-dependent blinking due to transitions towards excited (S0, S1) and dark states (S1, S2). **(b)** First, the initial model of the emitters’ positions is created by placing random seeds over the analyzed region, where each seed represents a fluorophore. **(c)** For each seed placed in the model, the behavior along the time is sampled based on the transition matrix and the initial probability as a Markov Chain Monte Carlo sampling. Local decisions are made by adding and removing one fluorophore at a time from the model. Using Bayesian Inference, the probability that the model with M fluorophores (F_M_) generates the diffraction limited data is computed. Finally, the SR image is created by accumulating the quantized positions of the probability map to the nearest pixel in the high resolution density image, applying a blurring to each fluorophore. **(d)** The parallelized version of 3B allows for multi-core parallel processing of sub-regions (or ‘patches’) of the input image stack, which aims for reduced computation times. **(e)** Super-resolved micrography using the parallelized version of 3B (Hernández et al., 2016) of rotavirus viroplasm where the green and red channels correspond to the VP6 and NSP2 proteins, respectively. Scale bars: 2 µm, insets: 1 µm.

The 3B algorithm initializes by placing uniform random seeds, assuming that fluorophores have the same probability of appearance over the entire region of analysis (**Figure 5b**). In the following step, the model is computed for each added fluorophore, F, and it is compared with the given input data (stack of images), D. These seeds are removed or added in such a way that the proposed model approaches the input data (**Figure 5c**). A set of models are generated with a different number of fluorophores, to maximize the probability of the model compared to the input data.

In summary, the comparison is performed by adding or removing fluorophores in the entire region, making local decisions, and then the model is re-optimized until the algorithm is stopped. Model convergence is said to be achieved when there is no significant variance between reconstructions of consecutive iterations (200 iterations are recommended for model convergence) (Cox et al., 2011). The reconstructed super-resolution image is a probability map of the position of the fluorophores, where the intensity of the pixels depends on the probability that a fluorophore is located in the region of interest.

When the 3B analysis was released, the execution required a sufficient level of programming skills, nevertheless, an Image-J plug-in was developed (Cox et al., 2011; Rosten, Jones and Cox, 2013). Both the script and the plug-in have the drawback of the high computation time required to analyze a set of images. The Image-J plug-in still required 6 hours to analyze a 1.5 × 1.5 μm area with 200 frames in the stack (Cox et al., 2011; Rosten, Jones and Cox, 2013). Despite the extended compute time, the Image-J plug-in is user friendly, since it only requires information about the FWHM associated with the PSF, the pixel size (in nm) and the initial number of seeds (ROI width per height/10), which is an estimation of the number of emitters in the ROI (Region of Interest).

Subsequent developments of 3B aimed to reduce the computational cost, by parallelizing the analysis with methods such as cloud computing (Hu et al., 2013) and cluster computing, or in a conventional personal computer (**Figure 5d**) (Hernández et al., 2016). Since the computation time for 3B depends on the size of the images to be analyzed, one way to reduce the computational cost was to divide the size of the images to be analyzed by regions of smaller size, in this way, each region is analyzed separately. At the end, the SR image is generated by stitching the results of each SR region. As this process can result in artifacts in the SR image, a study of the optimal overlap between the ROIs was carried out to reduce these artifacts (Hernández et al., 2016).

A more recent development, Bayesian localization microscopy based on Fluorescence Intensity Distribution (FID3B) performs a statistical analysis to improve the accuracy of the initial seeds of the emitters based on the pixel intensity, reducing the computational time and improving the SR image (Xu et al., 2015). Later, Quick-3B was developed. It combines the *k*-means clustering algorithm and a modified 3B analysis with a limited forward algorithm, which accelerates computation time 17-fold compared to 3B (Xu et al., 2017). Also in this work, SIngle Molecule guided BAyesian localization microscopy (SIMBA) is presented as a way to reduce the discontinuous structures that the 3B method creates as artifacts (Xu et al., 2017). However, SIMBA requires two types of emission fluorescent signals, using an SMLM algorithm (PALM) as an initial guide for the Bayesian analysis. Recently, Live-SIMBA was published, a plug-in for ImageJ based on SIMBA which does not necessarily require a dual-channel dataset. It is characterized by a computation time reduced thousands of times compared with 3B and an acceleration 25-fold compared to SIMBA (Li et al., 2020).

In 2019, another variant of the original 3B algorithm (3B-ODE) was presented to improve the convergence of the model, and the accuracy for estimating the probability map of the fluorophore positions (Suárez et al., 2019). It consists in calculating the transition probabilities between states by fitting the experimental data with ordinary differential equations. 3B-ODE models and fits velocity constants for the electronic transition between on and off state experienced by the fluorophores within the diffraction-limited imaged stack (Suárez et al., 2019). The BAyesian Multiple-emitter Fitting (BAMF) is also based on a 3B algorithm, which uses Reversible Jump Markov Chain Monte Carlo (RJMCMC) combined with MCMC sampling. The advantage of BAMF is that it incorporates the photophysical information of the sample and the density of the emitters for the creation of the model as prior information, allowing the adjustment of multiple emitters and removing the heterogeneous background. This method also provides the uncertainties in the number of emitters and the locations of the most likely model (Fazel et al., 2019).

Finally, Radial Fluctuation Bayesian Analysis (RFBA), is a proposed method that uses light-sheet microscopy with Bessel plane illumination for 3D SR imaging (Chen et al., 2020). For this, the initial points of the 3B algorithm are calculated using Super-Resolution Radial Fluctuations (SRRF) and the model optimization is based on these same locations generated by SRRF, reducing the computation time to half of that required for 3B.

3B analysis is a powerful technique that can be used for live cell imaging of samples labeled with standard fluorescent proteins. In a wide-field experiment, relatively few frames (hundreds) are needed to reconstruct a super-resolution image. It can accomodate overlapping of fluorophores and achieves a resolution of ∼ 40 nm (Suárez et al., 2019). All current 3B analysis implementations require programming skills (except the ImageJ plug-in, canonical 3B and Parallel3B) and, in some cases, are no longer being maintained or there is no open-source implementation available. Overall, the main drawback of the 3B analysis is the high computation time. However, this problem has been addressed by improvements or optimizations in the algorithm, or by adjusting the parameters such as the number of initial seeds or the transition matrix for a faster model convergence.

#### **E**ntropy-based **S**uper-resolution **I**maging

At the time of the publication of Entropy-based Super-resolution Imaging (ESI) (Yahiatene et al., 2015), improved versions of cSOFI and 3B had been published that solved some problems from their original publications, like the consideration of non-linear brightness, blinking of the fluorophores and the computational time to analyze the input data (Geissbuehler et al., 2012; Hu et al., 2013).

ESI was the first FF-SRM method that required only 100 frames collected in a fluorescent scene to reconstruct a super-resolution image (Yahiatene et al., 2015). This is achieved by analyzing fluorescence fluctuations through the local information content, pixel-wise, of the temporal dynamics of the fluorescences stored in the image stack. The rationale of ESI is that, in a time-lapse stack, each pixel harbors the intensity fluctuation of a specific place in space. The information-based entropy depends on the frequency of occurrence of fluorescence events and, by doing so, a highly fluctuating source will have high entropy, while a constant signal (such as a non-blinking emitter) or a background pixel would be discarded (**Figure 6a**). If a fluorophore is found in a pixel, its temporal fluctuations will contain more information (entropy) than the background due to the photon emission process.

**Figure 6.**
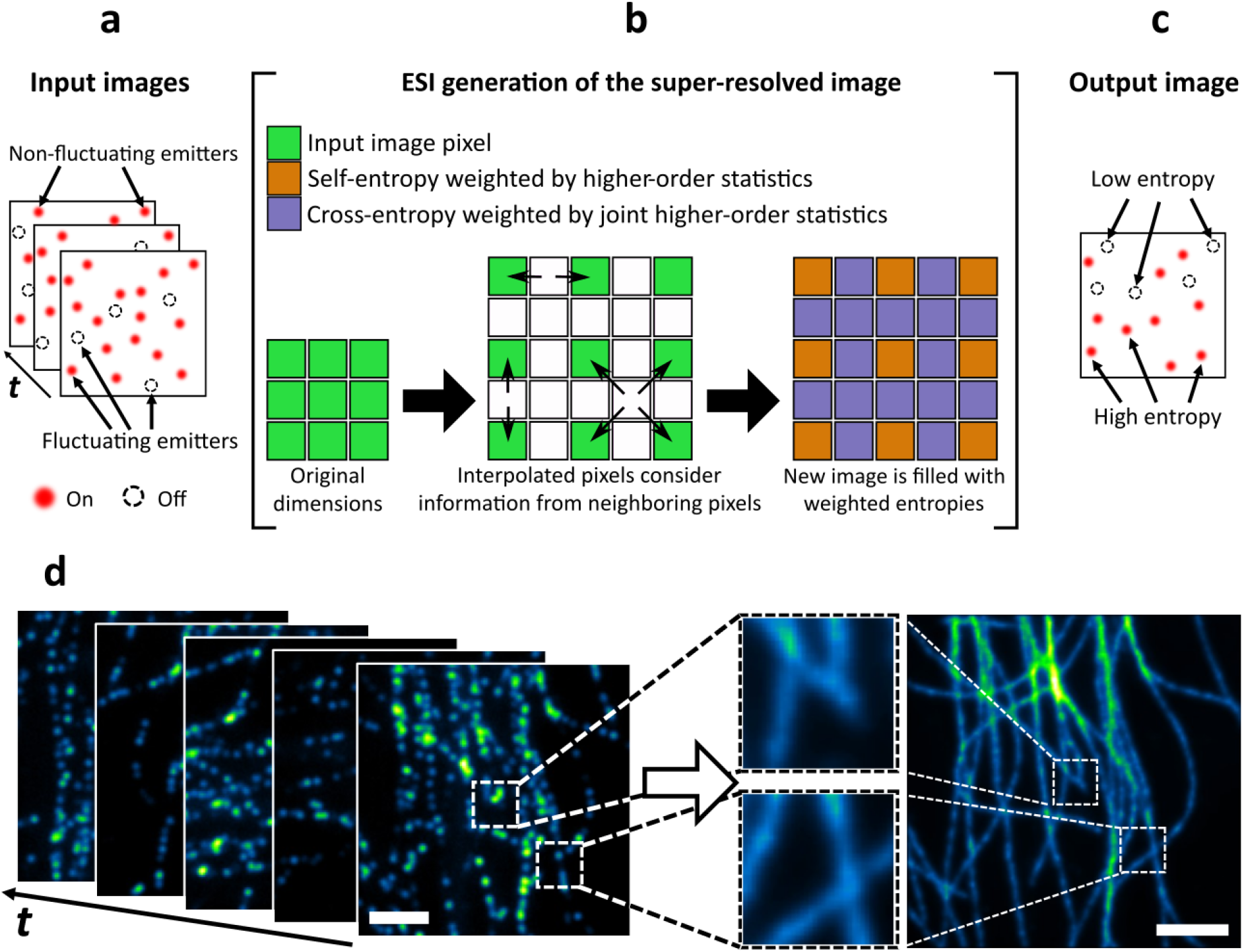
Entropy-based Super-resolution Imaging. **(a)** The input image stack consists of a set of fluctuating and non-fluctuating fluorophores. **(b)** The super-resolved ESI image is created by interpolation of the purple pixels in between the corresponding original green pixels. The orange pixels in the new image have a value that corresponds to the information content of the original image pixels (green), while the interpolating pixels (purple) consider the information of their neighbors. This scheme represents a single iteration; more iterations can be performed by using several output ESI images as new input. **(c)** The resulting image contains only the information of those emitters whose degree of variation (entropy) was high enough for detection. **(d)** Super-resolved micrography of simulated microtubules (Sage et al., 2015). Example generated using the ESI plugin for ImageJ with parameters: output images = 100, bins for entropy = 100, order = 1, multicore = enabled. Scale bars: 2 µm.

ESI, like SOFI and its variants, employs a pixel-wise calculation to extract the temporal information, with the difference that it calculates the information-based entropy (Yahiatene et al., 2015), instead of cumulants as in SOFI (Dertinger et al., 2009). While cumulants are an alternative of moments of a distribution (fluorescence intensities in this case), entropy measures the information content of the fluorescence signal. This entropy is weighed by the central moment (moments about the mean) of order n in the pixel; meaning that the central moment of order 2 is the pixel variance. Also, it is important to note that the 2nd and 3rd central moments are equal to the 2nd and 3rd cumulants, respectively, although at higher orders moments and cumulants generally differ.

A similarity between the ESI and SOFI is that ESI generates virtual pixels by exploiting the cross-entropy of neighboring pixels (combined with higher order statistics) (Yahiatene et al., 2015), and XC-SOFI generates extra pixels by computing the cross-cumulants of neighboring pixels (Dertinger et al., 2010). Since the cross-entropy is non-commutative, ESI calculates the average of the two possible cross entropies between 2 pixels. In the process of generating a new super-resolved image, ESI creates a magnified grid of the original image by interpolating one pixel in between each original pixel, as shown in **Figure 6b**.

Regarding their ability to reduce the PSF width (meaning enhancing resolution), while SOFI directly raises the PSF to the order n of the cumulant (thus achieving improvement of √n in resolution for cSOFI and of n for bSOFI) (Dertinger et al., 2009)(Geissbuehler et al., 2012), ESI considers 2n while obtaining the higher order statistic of the pixels, and the authors show that ESI improves the resolution by √(2n).

In ESI, the pixels in the new grid that correspond to the original image (green) are assigned the value of their entropy weighted by a higher order statistical measure selected by the user. This higher order statistic is the central moment about the mean, meaning that an order of n = 2 will yield the variance at that specific pixel. For the interpolated pixels, their new value is calculated from the cross-entropy of the neighboring pixels weighted by the joint centralized moment of those same neighboring pixels. Following the example of XC-SOFI, this cross-correlation between neighboring pixels yields a true signal in contrast to a typical interpolation on the final image (**Figure 6 c, d**)

ESI has been deployed as a plug-in for ImageJ, which generates a 2X magnification per iteration of the analysis (Yahiatene et al., 2015). In this implementation, each new iteration takes the ESI stack (output) and uses it as input, so ESI will calculate the entropy and cross entropy of this new input data. No more than 3 iterations are recommended due to the non-linear decrease in contrast, as well as information loss by pixel size reduction (Yahiatene et al., 2015). The plug-in also requires the number of images in the output which will define how the initial stack will be subdivided to generate the SR images.

Overall, ESI is an FF-SRM approach that can work with as few as 100 wide-field images to provide a reconstruction with narrower structures. The combination of orders and iterations remains user-dependent, which can be adjusted for achieving noise suppression, however, there is always the possibility that this suppression eliminates signals from fluorophores. In addition, the iterative process itself has a drawback since the plug-in is restricted to produce only a 2-fold increase in spatial sampling and analyze only a fraction of the frames per iteration when more than one image in the output is desired. Additional settings to aid the reconstruction had included the use of chip-based waveguides for alternative illumination set up (Diekmann et al., 2017).

#### **S**uper **R**esolution **R**adial **F**luctuation

Super Resolution Radial Fluctuation (SRRF) was developed in 2016 (Gustafsson et al., 2016), it consists of two main parts, a spatial analysis in which the algorithm generates a radiality map per each raw image, and a temporal analysis of each radiality map by higher-order temporal statistics to generate a single super-resolved image (**Figure 7 a - c**) (Gustafsson et al., 2016). SRRF is capable of reconstructing an SRM image with only 100 frames, achieving lateral resolution of ∼ 60 nm on TIRF, Confocal Laser Scanning Microscopy, wide-field and traction force microscopy data sets (Gustafsson et al., 2016; Culley, Tosheva, et al., 2018); (Stubb et al., 2020).

**Figure 7.**
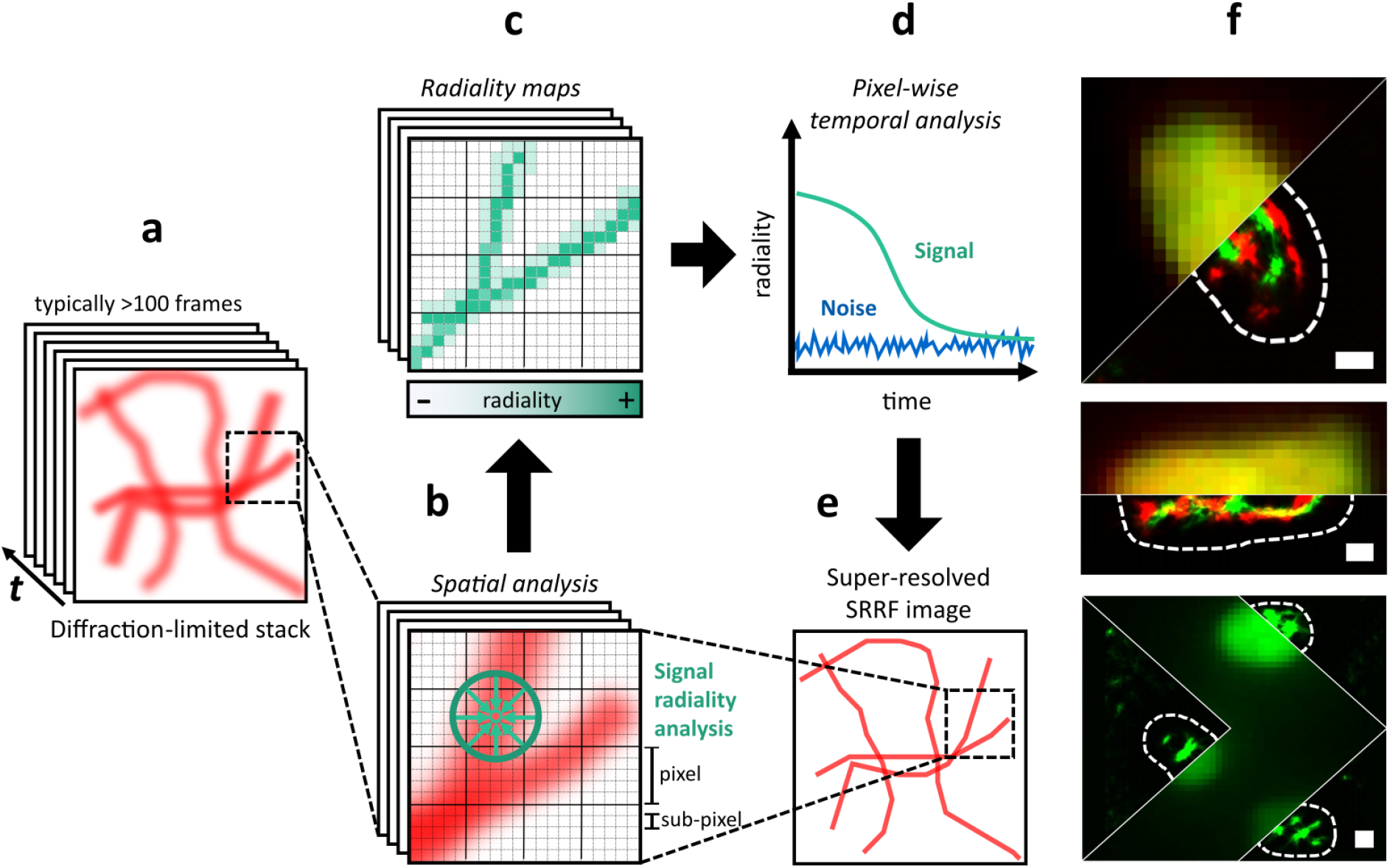
Super-Resolution Radial Fluctuations. (a) Each image of the input sequence is (b) sub-pixeled and on each of them a radiality measure is performed. (c) A radiality map is generated on every image from the sequence and finally, (d) they are correlated along time in order to (e) form the reconstructed image. (f) Super-resolved images of *E. coli* imaged on HILO microscopy, where the green and red channels correspond to mClover3 and mRuby3, respectively. Reconstructions were generated with the NanoJ SRRF plugin of ImageJ /FIJI with the parameters: Ring radius = 0.5, Magnification = 10, Axes in ring = 8. Scale bars: 350 nm.

The spatial analysis of SRRF begins with the generation of digital sub-pixels per each “real” pixel of the raw image sequence by a bicubic interpolation (**Figure 7b**). Next, each sub-pixel is assigned a value according to the probability that it has an emitting fluorophore. This value is generated by radiality maps (**Figure 7c**), which measure the degree of convergence of intensity gradient vectors on one sub-pixel basis. If the sub-pixel has a fluorophore, the convergence of the vectors will be higher than those coming from the image background.

The radiality maps can achieve better results for the SRM reconstruction by using a smaller ring radius and more axes in the ring. On the SRRF ImageJ plug-in (NanoJ-SRRF), the default ring radius is 0.5 pixels and the number of axes are six. It is important to consider that modification in these values will affect the computing time, the resolution achieved and the propensity to generate artifacts on the final SRM reconstruction (Gustafsson et al., 2016; Culley, Tosheva, et al., 2018). The maximal magnification factor achieved by NanoJ-SRRF is 10x, and it can be defined by the user according to their necessities.

Within a single radiality map, SRRF is capable of distinguishing two fluorophores separate by 0.7 the FWHM of the PSF, however, the noise of the image can be interpreted as a fluorophore signal and generate artifacts. Temporal analysis is needed to mitigate artifacts as the radiality peak on the background sub-pixels will be uncorrelated on time (**Figure 7d**). Like the other FF-SRM methods, the temporal analysis is affected by the sample movement thus, the NanoJ-SRRF plugin allows the application of a drift correction table, defined by the user or calculated by the algorithm.

SRRF encompasses any of 4 temporal analyses: Temporal Radiality Maximum (TRM), Temporal Radiality Average (TRA), Temporal Radiality Pairwise Product Mean (TRPPM), and Temporal Radiality Auto-Cumulant (TRAC) (Gustafsson et al., 2016). The temporal analysis method should be chosen depending on the acquired data; for example, TRA is recommended with noisy images and TRM is preferred when the images contain constantly emitting sources. Either TRA or TRM denoise the radiality map because they project the maximum and average values from the stack of the radiality maps. The TRPPM and TRAC orders 2, 3 and 4 (similar to those used in cSOFI) are higher order statistics methods that improve contrast, fidelity, and resolution on the SRM image (Gustafsson et al., 2016).

SRRF has been extensively utilized in bioimaging because it does not require the use of specific fluorophores, buffers or microscopes, and it only needs 100 frames to reconstruct a super-resolved image. It is compatible with live cell imaging since the use of high illumination power is unnecessary, which diminishes phototoxicity on the sample. SRRF has been used in combination with SIM (fFE-SIM) (Yao et al., 2021), STORM (Han et al., 2019), AiryScan (FEAST), and expansion microscopy with Airyscan (Ex-FEAST) (Wang et al., 2020) achieving resolutions of ∼ 32 nm, ∼40 nm and ∼26 nm respectively.

Unfortunately, SRRF is prone to generate artifacts on noisy images and very high-density fluorophore samples, and often over narrows the structures on the image (Yi et al., 2019; Pawlowska et al., 2022). Therefore, it is recommended to optimize the input parameters (especially ring radius, axes and temporal analysis) with error mapping approaches (Culley, Albrecht, et al., 2018).

An improved version has been released named enhanced SRRF (eSRRF) (Gustafsson et al., 2016; Laine et al., 2022), which improves the original algorithm by changing three main aspects; the first corresponds to the interpolation performed to generate sub-pixels in eSRRF - this process is performed by Fourier transform interpolation which minimizes the artifacts in the final SR image. Secondly, the radiality maps are now calculated by radial gradient convergence (RGC) which is calculated by a weighted factor map based on the Radius (R-value) defined by the user and the intensity gradient of each pixel on the original image. Finally, a new parameter defined as Sensitivity (S-value) is included which allows better control of the PSF sharpening performed by the RGC.

The eSRRF plug-in has become more user-friendly by adding a parameter sweep on which different values of the radius and sensitivity are compared using the error and mapping tool SQUIRREL, diminishing the probability of artifacts by a non-optimized selection of the algorithmic parameters. In general, low sensitivity and radius values increase image fidelity; in contrast, the resolution is increased with higher S values at the cost of lower image fidelity.

Currently, SRRF and eSRRF are available as ImageJ /FIJI plug-in NanoJ SRRF and NanoJ eSRRF, respectively (Gustafsson et al., 2016; Laine et al., 2022). In addition, SRRF has been deployed in python (Han et al., 2019), with a decrease of up to 78-folds the processing time (compared with SRRF) by allowing parallel computing supported by Compute Unified Device Architecture (CUDA) (code not available). SRRF can be performed in real-time with parallel GPU computing with SRRF-Stream and SRRF-Stream+, which are only available for microscopes with specific Andor and Sona cameras.

#### **Mu**ltiple **Si**gnal **C**lassification **Al**gorithm

Just a few months after SRRF was published back in 2016, the Multiple Signal Classification (MUSIC) Algorithm (MUSICAL) for FF-SRM was released, presented as an ideal tool for live-cell nanoscopy due to the minimum dataset size requirement: as few as 50 frames (**Figure 8a**) can provide a resolution enhancement, achieving ∼50 nm in live-cell experiments. Because of the low number of frames required, MUSICAL can reconstruct super-resolved images at a temporal spacing of less than 50 ms (with a maximum frame rate of 1000 fps), and represents a powerful option to study fast biological processes with nanoscopic resolution (Agarwal and Macháň, 2016).

**Figure 8.**
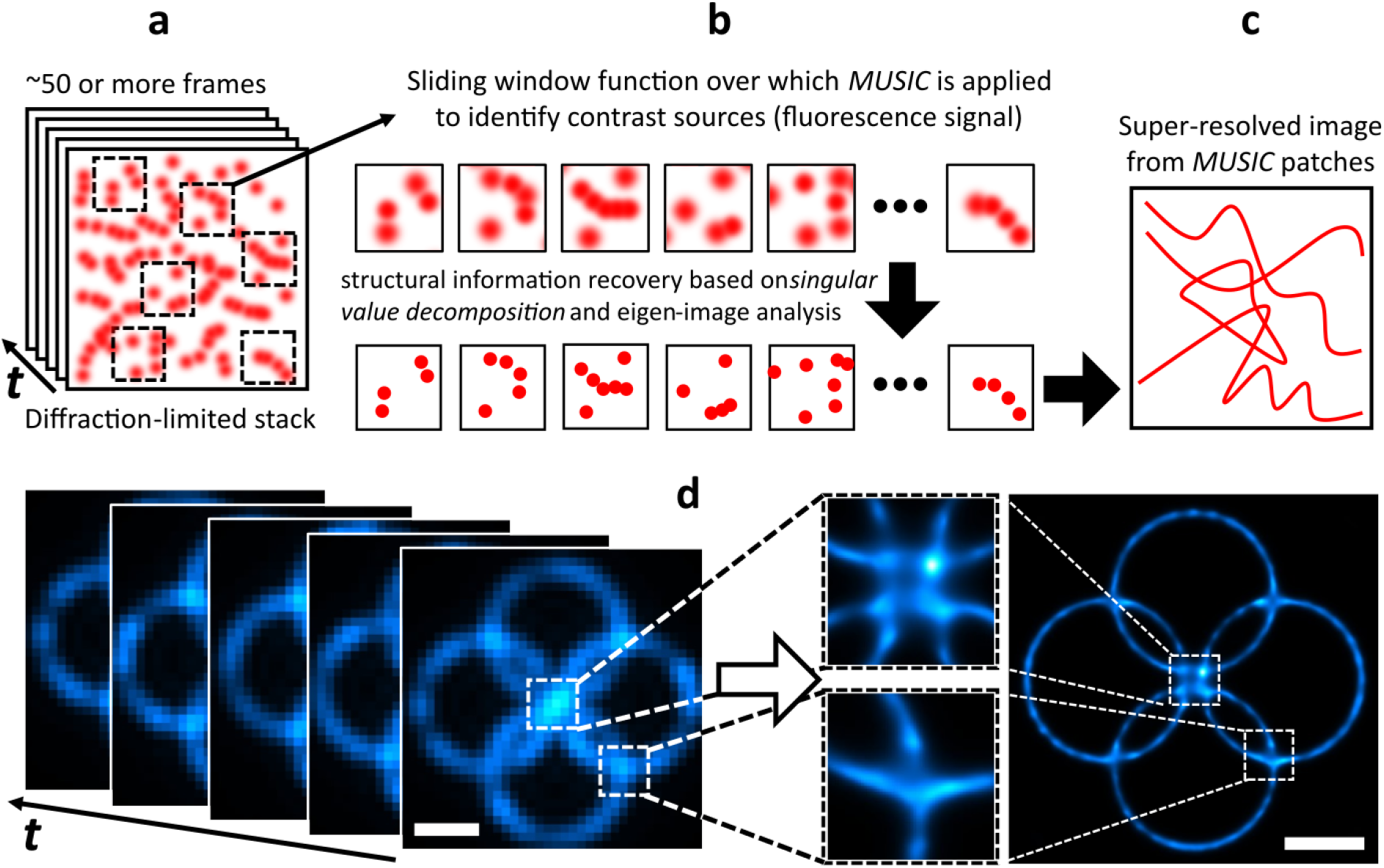
Multiple Signal Classification Algorithm. (a) For every pixel of the input image stack, a spatial window with a size which is approximate to the FWHM the microscope’s PSF (in pixels) is considered. This window is then scanned across the whole image. (b) This generates multiple temporal image ‘patches’, over each of which SVD is performed and the eigenimages that represent the distribution of fluorophores for each of these regions are calculated. Then, based on the signal and noise boundaries established for each image patch, structural information is recovered. Each pixel is then divided into a sub-pixel grid and the coordinates of the emitters are estimated. (c) Finally, all the sub-pixel MUSIC patches are stitched together and the super-resolved image is formed. (d) Super-resolved image of a synthetic nanoscopic structure with simulated blinking fluorophores placed along each ring (Agarwal and Macháň, 2016). Example generated using the MusiJ plugin for ImageJ with parameters: emission = 510 nm, numerical aperture = 1.49, magnification = 1, pixel size = 65 nm, threshold = -0.5, alpha = 2, subpixels per pixel = 5. Scale bars: 500 nm.

Based on the MUSIC (Multiple Signal Classification) approach (Schmidt, 1986), MUSICAL relies on the Singular Value Decomposition (SVD) principle. Briefly, SVD highlights the most prominent sources of variation within a system or a set of measurements by projecting the data onto a feature space (Stewart, 1993). Unlike MUSIC, MUSICAL overcomes the need for impractically large data sets by applying a sliding soft window in which the MUSIC algorithm is implemented (**Figure 8b**). Given that the number of images required by MUSIC scales proportionally to the number of contrast sources (fluorophores) that are present, this sliding window feature restrains computational demands while providing accurate signal reconstructions in terms of resolution and image quality (Agarwal and Macháň, 2016).

First, once the sliding window function is defined and the image patches are created (**Figure 8a**), MUSIC calculates their eigenimages and their corresponding eigenvalues in order to recover the most prominent features of the sample by identifying the contrast sources within the signal (fluorescence) (**Figure 8b**). Eigenimages are a collection of images that, summed together using their eigenvalue as weights, can be used to obtain the input image (Turk and Pentland, 1991; Leonardis and Bischof, 2000; Monwar, Rezaei and Prkachin, 2007). The SVD principle also helps to suppress noise by filtering-out those eigenimages with lower eigenvalues, for which a user-defined threshold is used. Small (or near-zero) eigenvalues are typically linked to a poor image signal and high noise. MUSICAL relies on the decomposition of the observation space (in this case, an image) into a source/signal (range) subspace and noise (null) subspace. This information helps the algorithm establish the boundaries of what is signal and what is background noise.

Next, structural information provided by MUSIC is enriched with prior knowledge of the PSF of the optical system, which provides regions of the likely position of the emitters by computing their projection in both the range and null subspaces. The algorithm determines if the PSF is statistically represented by each subspace by analyzing their corresponding eigenvalues. To improve the estimation of the emitter location when the PSF calibration accuracy is low, a parameter *alpha* is used to narrow down the expected spread of the emitter. Typically, this parameter is chosen to be *alpha ≥ 2*. Finally, using the user-defined parameter *sigma_0*, which serves as a threshold for the singular values of each eigenimage, the pixels in the image that correspond to the signal (range subspace) are separated from those that correspond to background noise (null subspace) (**Figure 8b**). This last step generates the super-resolved patches that are then stitched together to reconstruct the entire image (**Figure 8c**). An example of the processing of MUSICAL over simulated data is presented in **Figure 8d**.

Worth mentioning is that this patch (or sub-region) analysis is different from that carried out by the parallelized version of 3B, which physically assigns each data patch to a specific core of the CPU (**Figure 5d**) for parallel processing, with the only aim to reduce the processing time. For the case of MUSICAL, the use of patch-level analysis impacts directly on the quality of the reconstruction since the MUSIC approach performs differently as a function of image size and emitter density.

MUSICAL does not require special blinking-inducing photochemical treatment. It also performs relatively well in high-density fluorophore conditions (Agarwal and Macháň, 2016; Solomon et al., 2018). Unlike other methods such as STORM, which rely on long dark fluorophore states, MUSICAL performs well in both long- and short-term dark state conditions. Additionally, despite blinking behavior, the sliding window feature of MUSICAL allows the faithful reconstruction of the nanoscopic structures within the sample, while maintaining the required dataset size small enough (a minimum of ∼50 frames) (Agarwal and Macháň, 2016). This facilitates the study of dynamic biological phenomena. Although less computationally demanding, MUSICAL performs poorly as the SNR diminishes (Opstad et al., 2020). This condition becomes inconvenient in reduced light level experiments and makes this method prone to noise-related artifact generation. Also on MUSICAL, both reconstruction ability and precision decrease proportionally with the sample brightness, which means that a poor photon emission will likely lead to false negatives in the resulting MUSICAL super-resolution image. Increasing the laser power may help but it can lead to bleaching and phototoxicity. It is worth mentioning that MUSICAL requires a relatively accurate prior knowledge of the PSF of the microscope, which may otherwise lead to mild artifact generation. However, it behaves somewhat flexibly with respect to the estimation due to the *alpha* parameter mentioned above. The MUSICAL ImageJ plug-in includes an automatic PSF estimation tool based on optical parameters (emission wavelength, Numerical aperture, magnification and pixel size) (code available at https://github.com/sebsacuna/MusiJ).

Similar to MUSICAL, another SRM method that provides resolution enhancement based on eigenimages calculation is SCORE (Deng et al., 2014). This method relies only on the information related to the range subspace, which limits its ability to reduce background-induced artifacts. Also, SCORE lacks a sliding window function which, in combination with a cost-minimization iteration-like procedure, makes it a more computationally demanding method.

Overall, MUSICAL offers comparable and often superior results against most previously reported SRM methods, increasing resolution to below 50 nm. It performs competitively in a variety of experimental scenarios (such as fixed or live-cell imaging and 3D nanoscopy, for example) and also in terms of complexity, dataset size, computational times and highest resolution attainable. Its ability to reconstruct a super-resolved image from a dataset of as small as ∼ 50 frames makes it a great choice for live-cell nanoscopy. It is currently implemented in the MATLAB and FIJI/ImageJ platforms and does not require specialized instrumentation or complex sample treatment (Acuña et al., 2020). However, reconstruction quality is severely affected by factors related to the overall quality of the signal, which includes fluorophore brightness, blinking dynamics and SNR.

#### **M**ean-**S**hift **S**uper **R**esolution microscopy

The Mean-Shift Super Resolution (MSSR) microscopy approach can obtain a super-resolved image from a single diffraction-limited image, achieving a resolution improvement of about a half (up to ∼140 nm) (**Figure 9a**) (García et al., 2021). MSSR is based on an idea similar to the radiality maps in SRRF. In addition, MSSR can be used as an FF-SRM method by incorporating a temporal analysis, providing further spatial resolution enhancement, allowing the reliable discrimination of neighboring emitters separated by 40 nm, by means of analysing fewer than 30 frames (**Figure 9b**).

**Figure 9.**
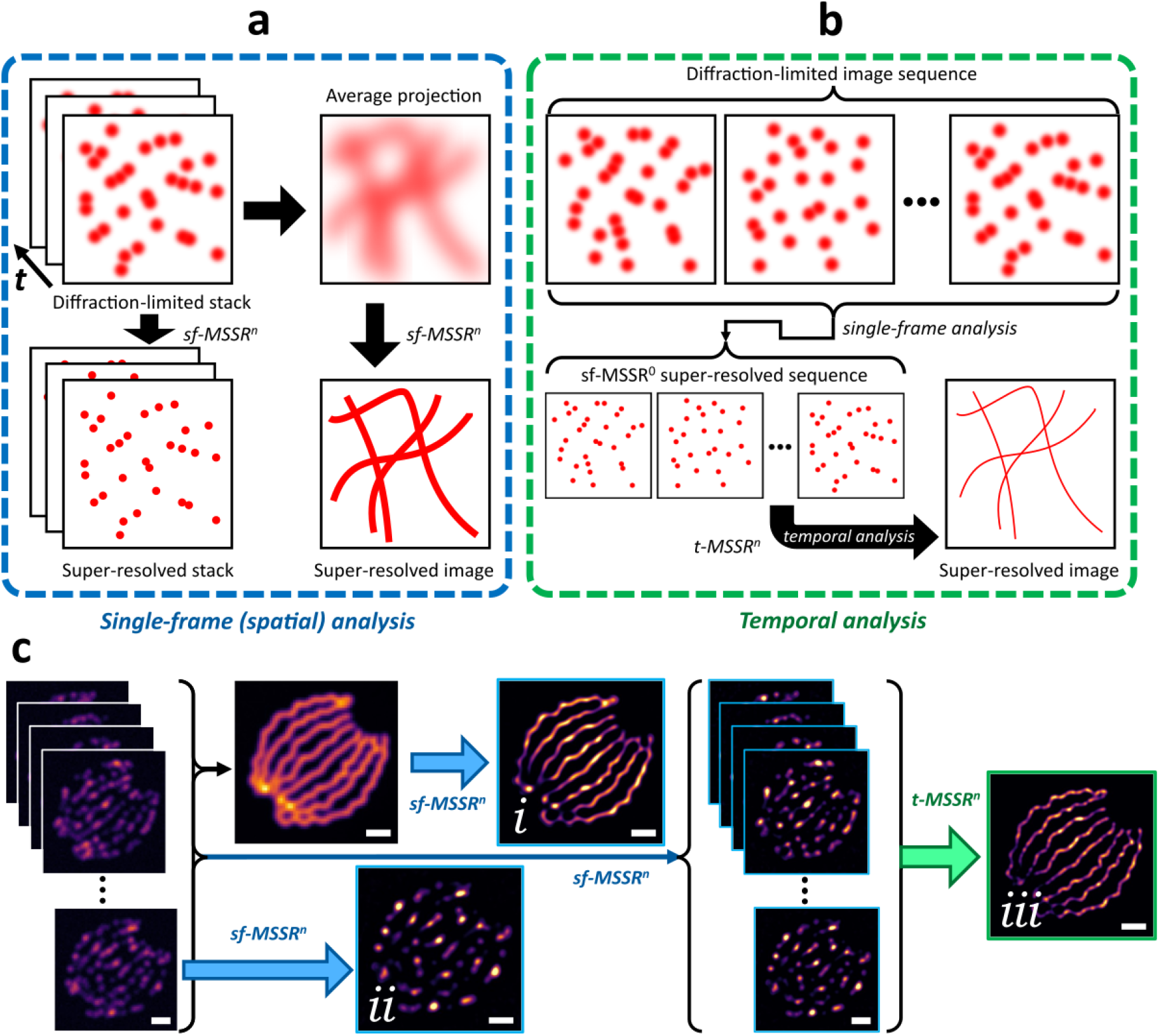
Mean-Shift Super Resolution microscopy. **(a)** Single-frame MSSR (sf-MSSR^n^) of a given order *n* reduces emitter width resulting in a super-resolved image. This can be applied to either each image of a diffraction-limited stack or to its average projection. **(b)** In MSSR temporal analysis (t-MSSR^n^), first, the super-resolved stack is obtained (a common step in both processing modalities), and then a given pixel-wise temporal function is used to generate the reconstruction. (c) sf-MSSR^n^ (*i & ii*) and t-MSSR^n^ (*iii*) result from MSSR applied on a simulated microtubules data set (Sage et al., 2015). Example generated using the MSSR plugin for ImageJ with parameters: Amp = 5, FWHM = 2, order = 1, interpolation = bicubic, meshing minimization = enabled, temporal analysis = variance. Scale bars: 1 µm.

MSSR analysis shows striking denoising capabilities that outperform other FF-SRM approaches, allowing the robust scrutiny of nanoscopic scales in a wide range of signal-to-noise ratio (SNR) conditions (García et al., 2021). MSSR is relatively robust in processing low- or high-density fluorophore images, and achieves comparable results in processing images collected with either CCD, sCMOS, or photomultiplier-based laser scanning technologies.

MSSR is based on MeanShift (MS) theory, which estimates local similarity properties between a central point and its neighbors (Fukunaga and Hostetler, 1975; Cheng, 1995). MS is a vector that lies on the second-order derivative space related to the data points and always points to the direction of maximum local density of information. MS was conceptualized to iteratively climb through data points until reaching local density modes (stationary points for the iterative procedure) in the data space (Comaniciu and Meer, 2002; Rao, Martins and Príncipe, 2009). Most MS applications are distinguished by two main features: the search type for local density modes and the application of iterative procedures to find these modes (Comaniciu and Meer, 2002; Barash and Comaniciu, 2004; Hu, Juan and Wang, 2008; Fazekas et al., 2021). These two features define the classical iterative procedure of MS. However, unlike the classical iterative procedure of MS, MSSR does not search modes along the data space and computes only the first MS value; this means that MSSR does not require MS iterations on its calculations (García et al., 2021).

The single-frame MSSR analysis (sf-MSSR^n^) (**Figure 9a**) is based on local kernel density estimation, hence, both spatial and range parameters are required (García et al., 2021). The spatial parameter is calculated from the optical properties of the imaging system and fluorophore features, namely, the pixel size of the diffraction-limited image, the numerical aperture of the imaging lens, and the emission wavelength of the fluorophore. The range parameter is defined automatically by the maximum difference of intensities on each neighborhood that slides over the image.

MSSR offers the ability to select an iterative approach that provides higher spatial resolution (MSSR^n^) (García et al., 2021). The authors refer to this as MSSR order, denoted by n. MSSR zero order (n=0) is constituted by the computation of MS, reducing the FWHM of isolated emitters by about half. MSSR of higher orders (integer n>0) is performed as an iterative procedure that applies basic algebraic functions, such as subtraction, multiplication, complement and normalization, on consecutive resulting images, which reduces even further the FWHM of emitters. MSSR provides a further improvement in resolution as the order increases.

The sf-MSSR^n^ resolution limit is reduced to 0.64 times the FWHM, and emitters at a smaller distance can not be distinguished by sf-MSSR analysis (for comparison, SRRF radiality maps analysis provides an improvement in resolution equivalent to 0.7 times the FWHM). In general, higher order MSSR analysis preserves the highest intensity of the original image, but decreases the lower intensities progressively. For this reason, it is suitable to remove noise but harmful to the quality of the reconstructed image. Authors recommend using MSSR orders not greater than 3.

The MSSR temporal analysis (t-MSSR^n^) integrates all the information over the sf-MSSR^n^ super-resolved stack by applying a pixel-wise temporal function (PTF) (**Figure 9b**). Each type of PTF has advantages depending on the nature of the fluorescence dynamics of the image stack to be processed. The t-MSSR^n^ analysis achieves a higher resolution corresponding to 0.21 times the FWHM.

Since MSSR is not limited by detector architecture and can process both single images and image stacks, there are several scenarios of fluorescence microscopy and bioimaging where MSSR offers good performance (**Figure 9c**). Among other microscopy fields, sf-MSSR^n^ is feasible on single-particle tracking or fixed-cell imaging microscopy, processing either a single image or each plane of a z-stack to obtain a 3D reconstruction. Given that sf-MSSR^n^ has the property to operate over a single image, it is easily combinable with all FF-SRM methods previously described to increase resolution on super resolved images. On the other hand, t-MSSR^n^ is suitable for temporal multi-frame analysis on fixed cells. In addition to the above, MSSR is compatible with other SRM techniques, such as SIM, STED, SOFI, 3B-ODE, ESI, MUSICAL and SRRF, allowing a further resolution enhancement when applied to their super-resolved images. This algorithm is available as a user-friendly Fiji/ImageJ plug-in (García et al., 2021).

### Artificial Intelligence-based FF-SRM techniques

Deep learning encompases a subset of machine learning algorithms, based on neural networks, which has gained popularity in recent years due to its great performance in different tasks such as segmentation (Ronneberger, Fischer and Brox, 2015), denoising (Krull, Buchholz and Jug, 2019), and SRM (Li et al., 2018; Nehme et al., 2018; Ouyang et al., 2018; Dardikman-Yoffe and Eldar, 2020; Chen et al., 2021). In supervised deep learning, a training dataset, consisting of the input image and the expected result (ground-truth), is used to tune the weights of a neural network. In the case of FF-SRM, the training dataset encompasses a collection of image pairs, consisting of a low-resolution (diffraction-limited) image, and its corresponding SR image (the ground-truth) (**Figure 10**).

**Figure 10.**
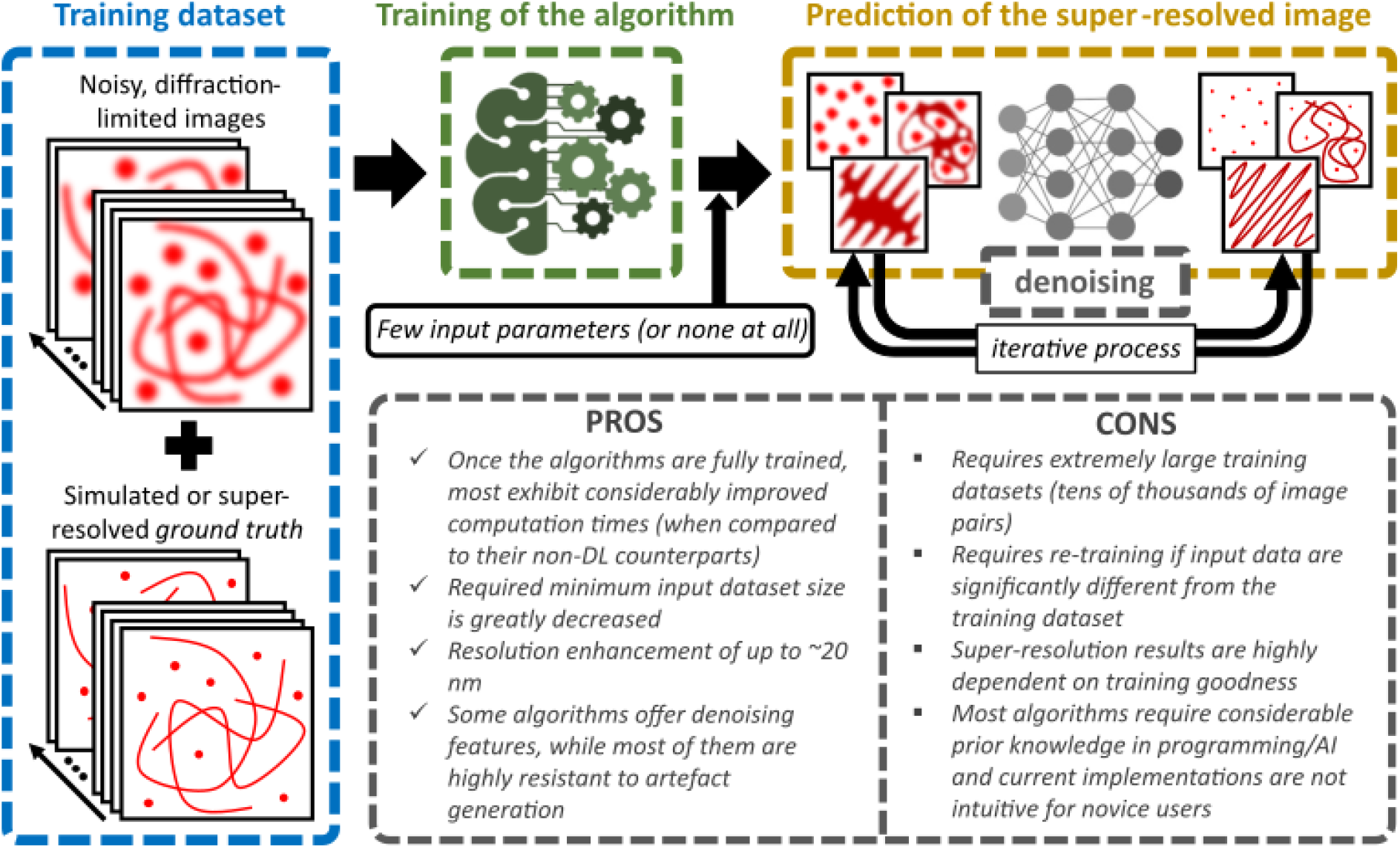
Deep-Learning Super-Resolution. Most deep learning algorithms require training in the form of thousands of diffraction limited images and their corresponding expected results of enhanced resolution and contrast. These image pairs are commonly simulated but can also be acquired in a real optical system and super-resolved with other SRM methods prior to training of the deep learning network. The demand of impractically large training data sets by deep learning approaches is compensated in the form of greatly improved computation times and almost parameter-free operation. However, caution must be taken as reconstruction quality is highly dependent on training fitness. Some of the most remarkable advantages and disadvantages of choosing deep learning approaches are displayed.

These approaches require a large quantity of images as training examples. For instance, Deep Learning guided Bayesian Inference (DLBI) is trained with 12,000 examples (Li et al., 2018), Deep-Storm with 10,000 examples (Nehme et al., 2018), and Single-Frame Super-Resolution Microscopy (SFSRM) with 1,000 examples (Chen et al., 2021). Such a large amount of training data is not always available to generate a good deep learning model, and there is not a general rule to decide the minimum number of images sufficient for the training.

DLBI (Li et al., 2018) is an algorithm that predicts an SRM image from a time series of high-density fluorescent images. In the first step, the time series images are transformed into an SRM image using a deep learning architecture called Generative Adversarial Network (GAN) (Goodfellow et al., 2014). The GAN is trained to simulate diffraction-limited stacks from a high-resolution image (12,000 images collected from Laplace-filtered natural images and sketches). Therefore, to simulate the training dataset DLBI encompases an algorithm which given a high-resolution image simulates the corresponding fluorescent time series images, by means of using a Markov model that switches between emitting, not emitting, and bleached states.

The algorithm takes into account expected photophysical properties of the emitters, such as switching probability between bright and dark excited states, as well as the PSF of the imaging system, in order to generate diffraction-limited images very similar to those from microscope acquisitions. In a second step, the high-resolution image predicted by the GAN is used to generate a set of possible fluorophore localization. Bayesian inference with the time series images is used to discard a given localization that is not well represented in the input dataset following a similar approach of 3B analysis (Cox et al., 2011).

An advantage of DLBI, compared to 3B analysis, is that it reduces the computational time up to 100 times and generates comparable quality SRM images, scored by using the metrics of peak signal-to-noise ratio, structural similarity, and SQUIRREL (Culley, Albrecht, et al., 2018). Even though DLBI has advantages over 3B and the results look promising for a wide range of data sets, its widespread usage as FF-SRM method is limited, since there is not an easy-to-use distribution/program. The authors released a GitHub repository (https://github.com/liyu95/DLBI), however, its use requires advanced programming skills.

SPArsity-based Super-resolution COrrelation Microscopy (SPARCOM) is an FF-SRM method that analyzes a temporal sequence of diffraction-limited images, which is achieved by taking into account the temporal fluctuation of the fluorescence emitters harbored within the collected images, their sparsity, and the fact that emitters are uncorrelated over time, aiming to model its behavior and to create a SRM image (Solomon et al., 2018). SPARCOM is based on the theory of SMLM and aims to recover the position of the emitters in a finer grid. Using prior information about the PSF of the imaging system, SPARCOM formulates a Multiple Measurement Vector (MMV) model, which is used to recover a SRM image (Davies and Eldar, 2012). In SPARCOM, the MMV model includes the use of a covariance matrix for the optimization of N^4^ variables (unknowns). Taking into account the fact that emitters are uncorrelated over time and sparse in the spatial domain, the number of unknown variables is reduced to N^2^, and these can be found using an Iterative Shrinkage-Thresholding Algorithm (ISTA) (Rozell et al., 2008).

SPARCOM can achieve a spatial resolution of ∼ 40 nm (similar to PALM and STORM), but through the analysis of only 50 frames. A disadvantage is the requirement of the prior-knowledge of the PSF, and the need for the regularization parameter ISTA (determined heuristically). The Learned SPARCOM (LSPARCOM) approach encompases an iterative algorithm which uses deep learning to predict super-resolution images without the need of making any assumption about the PSF of the imaging system (Dardikman-Yoffe and Eldar, 2020). It does not require fine-tuning of the optimization parameters as with SPARCOM, and produces similar or better SRM images than SPARCOM using the signal-to-noise ratio metric and visually.

In LSPARCOM, the PSF and the regularization parameter are learned from training data using a neural network approach (deep learning unfolding) (Karol Gregor and Yann LeCun, 2010). The training dataset is generated by setting the position of the emitters (ground-truth SRM image) from simulated biological microtubules or tubulins, and then simulating the diffraction limited images using microscopy parameters such as sample thickness, random activation, laser power, lifetime of fluorophores, noise, PSF size, amongst others, in order to generate diffraction limited images very similar to those from microscope acquisitions. The training dataset allows LSPARCOM to map simulated diffraction-limited images to SRM images, since simulated images are similar to experimental images. Furthermore, it also allows the mapping of experimental diffraction-limited images to the SRM reconstruction.

LSPARCOM generates SRM reconstructions similar to SPARCOM with a 5-fold computational time improvement (Dardikman-Yoffe and Eldar, 2020). In addition, LSPARCOM can generate good quality SRM images with as few as 25 images. A disadvantage of the LSPARCOM approach is the requirement for test low-resolution images to have a similar or not significantly different PSF from the images in the training dataset. If this is not the case, it can produce artifacts such as lines reconstructed as chains or intersection areas reconstructed as arcs. Similarly, LSPARCOM is sensitive to the pixel size of the simulated diffraction-limited images used during training (a pixel size of 100 nm was used during training). If a significantly higher or lower pixel size is used during the acquisition of the diffraction-limited image, it can generate inaccurate SRM images. Therefore, in order to have good quality results on a dataset with different PSF or different pixel size than 100 nm, LSPARCOM must be retrained, which as stated previously is not an easy task. The authors provide a link to the full code and a graphic user interface.

Recently, a novel algorithm based on Deep Learning, Single-Frame Super-Resolution Microscopy (SFSRM) (Chen et al., 2021), has been shown to achieve ∼20nm spatial resolution (10X resolution improvement compared to diffraction limited images). SFSRM uses two deep neural networks to obtain the SRM reconstruction. The first deep learning network, the Signal-Enhancement Network (SEN) receives as input the diffraction-limited image (low-SNR) and generates an image with High-SNR (noise reduced) while maintaining the same resolution. The second deep learning network, Super-Resolution Network (SRN) receives as input the High-SNR image and generates the SRM image. The training dataset for the SEN network (low-SNR and high-SNR) is obtained from fixed cells at different illumination intensities and from different microscopes (epifluorescence, TIRF, HILO, and confocal). This approach allows the network to reduce the noise in different microscopy techniques. The SRN network is then trained using simulated low-resolution images (100 nm pixel size) with their corresponding 10X high-resolution image (10 nm pixel size), and experimental diffraction-limited wide-field images with their high-resolution images, which are reconstructed using STochastic Optical Reconstruction Microscopy (STORM) (Rust, Bates and Zhuang, 2006).

The main feature of SFSRM is achieving ∼ 20 nm of resolution with a single diffraction-limited image. SFSRM was used to visualize cargo transport dynamics in a dense microtubule network. The highest resolution reached by SFSRM is limited by the resolution from the training SR images (obtained with STORM), hence, the resolution can be increased if SRM with higher resolution are used for the training (e.g. MINFLUX microscope). The main disadvantage is that the SFSRM depends on the patterns observed during the training dataset. Therefore, it can produce erroneous SRM reconstructions for input images with different topological structures than those used in the training set (necessitating the retraining of the SFSRM network). Currently, there is no source code available to test SFSRM.

Summarizing, deep learning based algorithms are a powerful approach to obtain SR images, since they are outperforming non-AI based approaches. However, their utilization is limited to a community with knowledge in artificial intelligence, due to the difficulty to train the networks which require to optimize the parameters of the network, programming skills, a large amount of training examples and high-computational resources (access to GPU). Efforts have been done to make deep learning algorithm available to novice users (Chamier et al., 2021; Gómez-de-Mariscal et al., 2021; Körber, 2022), for instance using ZeroCostDL4ML which provides easy-to-use Jupyter Notebooks for the training of deep neural networks in different tasks such as segmentation, denoising, object detection, super-resolution, etc. However, the only available method for super-resolution is Deep-STORM (Nehme et al., 2018). The research community must continue in the effort to make its algorithms easily available, and user-friendly, since typically the end-user does not have experience in artificial intelligence to train or use deep learning networks.

### SOFTWARE AVAILABILITY AND HOW TO CHOOSE THEM

Most of the methods presented in this work are implemented as free access software, however, the execution may require the use of multiple platforms such as Matlab, python, R, FIJI Plug-in, amongst others (Dertinger et al., 2009; Cox et al., 2011; Yahiatene et al., 2015; Agarwal and Macháň, 2016; Gustafsson et al., 2016; Li et al., 2018; García et al., 2021). Due to the constant improvement of computing, memory and storage capacity, it is possible to mention that most modern computers meet the requirements to use the SRM software developed so far, however, a GPU is recommended to deploy its full potential, speeding up the data processing.

There is a wide range of bioimaging and fluorescence microscopy applications in which FF-SRM methods can be used such as: immunofluorescence of fixed cells, live-cell imaging using organic dyes, fluorescent proteins or quantum dots, single-particle tracking, colocalization, 3D imaging, etc (Dertinger et al., 2009; Cox et al., 2011; Yahiatene et al., 2015; Agarwal and Macháň, 2016; Gustafsson et al., 2016; Li et al., 2018; García et al., 2021). When it comes to choosing new approaches to analyze data, some of the most important factors considered in the choice of a certain method are related to the overall cost-benefit of its use. In the context of SRM, the highest attainable resolution and the number of images required to achieve such goal are two closely related parameters that often lead to the selection of an approach over another.

Moreover, the nature of the input data themselves must be thrown into the equation as well to obtain the best results based on the available experimental conditions and resources. Methods such as SRRF, MUSICAL and MSSR are largely independent of fluorophore blinking dynamics and operate with relatively small dataset sizes, which makes these methods suitable for live-cell imaging (Agarwal and Macháň, 2016; Gustafsson et al., 2016; García et al., 2021). On the other hand, software availability and ease-of-installation are factors that might ultimately define the course of action. While most SRM approaches are implemented on many of the most popular image analysis platforms, others might not be so readily reachable. **Figure 3b** shows how all of the above-mentioned aspects compare among the SRM methods discussed in this review, according to the experience of each author; therefore, we recommend identifying cases in which a particular FF-SRM method has been used with similar purposes and experimental setups.

No matter which FF-SRM method is chosen, we recommend corroborating the SRM reconstruction with error mapping software like SQUIRREL (Culley, Albrecht, et al., 2018) or HAWKMAN (Marsh et al., 2021). Computing global resolution indexes like Resolution Scaled Pearson (RSP), Resolution Scaled Error (RSE) (Culley, Albrecht, et al., 2018) or Fourier Ring Correlation (FRC) (Banterle et al., 2013) are recommended to identify possible artifacts by loss of signal. With these metrics, a fair and systematic comparison between the FF-SRM methods could be achieved and misinterpretation or misuse of the software can be diminished.

## CONCLUDING REMARKS AND PERSPECTIVES

The development of FF-SRM methods have facilitated the temporally-resolved study of biology at the spatial nanoscale. Input images for most of these methods can be generated using microscopy platforms of general access, common fluorophores and simple sample preparation, making them suitable for use in most life science laboratories. That being said, it is nonetheless crucial to understand the basic concepts of these techniques and how each parameter will affect the final SRM reconstruction before choosing and employing a particular FF-SRM method.

Any FF-SRM technique is universally favored in all experimental scenarios. As is so often the case with microscopy, high performance of a particular technique by one criterion (i.e., highest resolution possible) may come at the expense of poorer performance in others (i.e., susceptibility for generating artifacts, slow computational performance). While we have striven to indicate the general performance (and trade-off) characteristics of the different approaches, there will always be scenarios in which any single algorithm may unexpectedly outperform another. This may be due to any combination of factors, such as a previously untried application case and the shifting nature of bottle-necks due to technological advancement over time. Due to the extensive circumstantial variables at play in each set of experiments, local factors will demand a degree of commitment to an empirical approach when choosing an FF-SRM technique for any given analysis task.

In this regard, the diverse range of options for FF-SRM mirrors the situation in the early (and still ongoing) years of the development of algorithms and implementations for SMLM image generation. A surge in interest in SMLM led to a rapid expansion of mathematical approaches for localizing single-molecule intensity fluctuations in temporal image series. Guidance for determining the strengths and weaknesses of the SMLM approaches advanced once canonical test data, simulated and real, were made available to the community and controlled criteria for assessing performance were applied in “challenges”. Most notably, the Single Molecule Localization Challenge (SMLC) (Sage et al., 2019) is available at https://srm.epfl.ch/. This “challenge” approach has also been adopted in the 3D Deconvolution Microscopy Challenge (3DDMC) at http://bigwww.epfl.ch/deconvolution/challenge/index.html.

An equivalent “challenge” for FF-SRM is yet to be devised and given the wide flexibility and applicability of FF-SRM techniques to a diverse range of microscopy modalities and experimental designs. A suitable bid for this dutty will be to assemble a canonical data set and universal performance criteria that will fairly and rigorously test the relative performance among the growing FF-SRM family members. The value of such a “challenge” is nevertheless unquestionable, in terms of time saved and errors avoided, for current and future researchers who seek to unlock the power of imaging to examine the most fundamental questions of biology at the highest spatial and temporal scales possible.

## CONTRIBUTIONS

A.A., E.B.A., and A.G. designed the review content: Introduction (A.A, E.B, E.T.), SOFI (E.B.A.), 3B (H.O.H), ESI (D.M.) SRRF (A.A.), MUSICAL (A.L.), MSSR (E.T.), Artificial Intelligence (P.H.) Software availability (R.P.). RD’A, C.W. and A.G. provided guidance, suggestions, concept and redaction validations. All the authors contributed to the conclusion section and approved the final version of the manuscript.

## ACKNOWLEDGMENTS

We thank Berenice Mora, David Torres, Sergio A. Valdés, Victor Abonza and Diana Velázquez for their valuable comments and suggestions after reading the manuscript.

## REFERENCES

Agarwal, K. and Macháň, R. (2016) “Multiple signal classification algorithm for super-resolution fluorescence microscopy,” Nature Communications, 7(1), p. 13752. doi:10.1038/ncomms13752.

Balzarotti, F. et al. (2017) “Nanometer resolution imaging and tracking of fluorescent molecules with minimal photon fluxes,” Science, 355(6325), pp. 606–612. doi:10.1126/science.aak9913.

Banterle, N. et al. (2013) “Fourier ring correlation as a resolution criterion for super-resolution microscopy,” Journal of Structural Biology, 183(3), pp. 363–367. doi:10.1016/j.jsb.2013.05.004.

Barash, D. and Comaniciu, D. (2004) “A common framework for nonlinear diffusion, adaptive smoothing, bilateral filtering and mean shift,” Image and Vision Computing, 22(1), pp. 73–81. doi:10.1016/j.imavis.2003.08.005.

Betzig, E. et al. (2006) “Imaging Intracellular Fluorescent Proteins at Nanometer Resolution,” Science, 313(5793), pp. 1642–1645. doi:10.1126/science.1127344.

Chamier, L. von et al. (2021) “Democratising deep learning for microscopy with ZeroCostDL4Mic,” Nature Communications, 12(1), p. 2276. doi:10.1038/s41467-021-22518-0.

Chen, R. et al. (2020) “Efficient super-resolution volumetric imaging by radial fluctuation Bayesian analysis light-sheet microscopy,” Journal of Biophotonics, 13(8). doi:10.1002/jbio.201960242.

Chen, R. et al. (2021) “Deep-Learning Super-Resolution Microscopy Reveals Nanometer-Scale Intracellular Dynamics at the Millisecond Temporal Resolution,” bioRxiv, p. 2021.10.08.463746. doi:10.1101/2021.10.08.463746.

Cheng, Y. (1995) “Mean shift, mode seeking, and clustering,” IEEE Transactions on Pattern Analysis and Machine Intelligence, 17(8), pp. 790–799. doi:10.1109/34.400568.

Comaniciu, D. and Meer, P. (2002) “Mean shift: a robust approach toward feature space analysis,” IEEE Transactions on Pattern Analysis and Machine Intelligence, 24(5), pp. 603–619. doi:10.1109/34.1000236.

Cox, S. et al. (2011) “Bayesian localization microscopy reveals nanoscale podosome dynamics,” Nature Methods, 9(2), pp. 195–200. doi:10.1038/nmeth.1812.

Culley, S., Albrecht, D., et al. (2018) “NanoJ-SQUIRREL: quantitative mapping and minimisation of super-resolution optical imaging artefacts,” Nature methods, 15(4), pp. 263–266. doi:10.1038/nmeth.4605.

Culley, S., Tosheva, K.L., et al. (2018) “SRRF: Universal live-cell super-resolution microscopy,” The International Journal of Biochemistry & Cell Biology, 101, pp. 74–79. doi:10.1016/j.biocel.2018.05.014.

Dardikman-Yoffe, G. and Eldar, Y.C. (2020) “Learned SPARCOM: unfolded deep super-resolution microscopy,” Optics Express, 28(19), p. 27736. doi:10.1364/oe.401925.

Davies, M.E. and Eldar, Y.C. (2012) “Rank Awareness in Joint Sparse Recovery,” IEEE Transactions on Information Theory, 58(2), pp. 1135–1146. doi:10.1109/tit.2011.2173722.

Dedecker, P., Duwé, S., et al. (2012) “Localizer: fast, accurate, open-source, and modular software package for superresolution microscopy,” Journal of Biomedical Optics, 17(12), pp. 126008–126008. doi:10.1117/1.jbo.17.12.126008.

Dedecker, P., Mo, G.C.H., et al. (2012) “Widely accessible method for superresolution fluorescence imaging of living systems,” Proceedings of the National Academy of Sciences, 109(27), pp. 10909–10914. doi:10.1073/pnas.1204917109.

Deng, Y. et al. (2014) “Spatial Covariance Reconstructive (SCORE) Super-Resolution Fluorescence Microscopy,” PLoS ONE, 9(4), p. e94807. doi:10.1371/journal.pone.0094807.

Dertinger, T. et al. (2009) “Fast, background-free, 3D super-resolution optical fluctuation imaging (SOFI),” Proceedings of the National Academy of Sciences, 106(52), pp. 22287–22292. doi:10.1073/pnas.0907866106.

Dertinger, T. et al. (2010) “Achieving increased resolution and more pixels with Superresolution Optical Fluctuation Imaging (SOFI),” Optics Express, 18(18), pp. 18875–18885. doi:10.1364/oe.18.018875.

Dertinger, T. et al. (2012) “SOFI-based 3D superresolution sectioning with a widefield microscope,” Optical Nanoscopy, 1(1), p. 2. doi:10.1186/2192-2853-1-2.

Deschout, H. et al. (2016) “Complementarity of PALM and SOFI for super-resolution live-cell imaging of focal adhesions,” Nature Communications, 7(1), p. 13693. doi:10.1038/ncomms13693.

Diekmann, R. et al. (2017) “Chip-based wide field-of-view nanoscopy,” Nature Photonics, 11(5), pp. 322–328. doi:10.1038/nphoton.2017.55.

Fazekas, F.J. et al. (2021) “A Mean Shift Algorithm for Drift Correction in Localization Microscopy,” Biophysical Reports, 1(1), p. 100008. doi:10.1016/j.bpr.2021.100008.

Fazel, M. et al. (2019) “Bayesian Multiple Emitter Fitting using Reversible Jump Markov Chain Monte Carlo,” Scientific Reports, 9(1), p. 13791. doi:10.1038/s41598-019-50232-x.

Fukunaga, K. and Hostetler, L. (1975) “The estimation of the gradient of a density function, with applications in pattern recognition,” IEEE Transactions on Information Theory, 21(1), pp. 32–40. doi:10.1109/tit.1975.1055330.

Gallina, M.E. et al. (2013) “Resolving the spatial relationship between intracellular components by dual color super resolution optical fluctuations imaging (SOFI),” Optical Nanoscopy, 2(1), p. 2. doi:10.1186/2192-2853-2-2.

García, E.T. et al. (2021) “Nanoscopic resolution within a single imaging frame,” bioRxiv, p. 2021.10.17.464398. doi:10.1101/2021.10.17.464398.

Geissbuehler, S. et al. (2012) “Mapping molecular statistics with balanced super-resolution optical fluctuation imaging (bSOFI),” Optical Nanoscopy, 1(1), p. 4. doi:10.1186/2192-2853-1-4.

Geissbuehler, S. et al. (2014) “Live-cell multiplane three-dimensional super-resolution optical fluctuation imaging,” Nature Communications, 5(1), p. 5830. doi:10.1038/ncomms6830.

Geissbuehler, S., Dellagiacoma, C. and Lasser, T. (2011) “Comparison between SOFI and STORM,” Biomedical Optics Express, 2(3), pp. 408–420. doi:10.1364/boe.2.000408.

Gómez-de-Mariscal, E. et al. (2021) “DeepImageJ: A user-friendly environment to run deep learning models in ImageJ,” Nature Methods, 18(10), pp. 1192–1195. doi:10.1038/s41592-021-01262-9.

Goodfellow, I.J. et al. (2014) “Generative Adversarial Networks,” arXiv [Preprint].

Gustafsson, M.G.L. (2000) “Surpassing the lateral resolution limit by a factor of two using structured illumination microscopy,” Journal of Microscopy, 198(2), pp. 82–87. doi:10.1046/j.1365-2818.2000.00710.x.

Gustafsson, N. et al. (2016) “Fast live-cell conventional fluorophore nanoscopy with ImageJ through super-resolution radial fluctuations,” Nature Communications, 7(1), p. 12471. doi:10.1038/ncomms12471.

Han, Y. et al. (2019) “Ultra-fast, universal super-resolution radial fluctuations (SRRF) algorithm for live-cell super-resolution microscopy,” Optics Express, 27(26), p. 38337. doi:10.1364/oe.27.038337.

Hell, S.W. and Wichmann, J. (1994) “Breaking the diffraction resolution limit by stimulated emission: stimulated-emission-depletion fluorescence microscopy,” Optics Letters, 19(11), p. 780. doi:10.1364/ol.19.000780.

Hernández, H.O. et al. (2016) “High Performance Computer Applications,” Communications in Computer and Information Science, pp. 356–366. doi:10.1007/978-3-319-32243-8_25.

Hertel, F. et al. (2016) “RefSOFI for Mapping Nanoscale Organization of Protein-Protein Interactions in Living Cells,” Cell Reports, 14(2), pp. 390–400. doi:10.1016/j.celrep.2015.12.036.

Hofmann, M. et al. (2005) “Breaking the diffraction barrier in fluorescence microscopy at low light intensities by using reversibly photoswitchable proteins,” Proceedings of the National Academy of Sciences, 102(49), pp. 17565–17569. doi:10.1073/pnas.0506010102.

Hu, J.-S., Juan, C.-W. and Wang, J.-J. (2008) “A spatial-color mean-shift object tracking algorithm with scale and orientation estimation,” Pattern Recognition Letters, 29(16), pp. 2165–2173. doi:10.1016/j.patrec.2008.08.007.

Hu, Y.S. et al. (2013) “Accelerating 3B single-molecule super-resolution microscopy with cloud computing,” Nature Methods, 10(2), pp. 96–97. doi:10.1038/nmeth.2335.

Jacquemet, G. et al. (2020) “The cell biologist’s guide to super-resolution microscopy,” Journal of Cell Science, 133(11), p. jcs240713. doi:10.1242/jcs.240713.

Klar, T.A. et al. (2000) “Fluorescence microscopy with diffraction resolution barrier broken by stimulated emission,” Proceedings of the National Academy of Sciences, 97(15), pp. 8206–8210. doi:10.1073/pnas.97.15.8206.

Kodama, Y. and Hu, C.-D. (2012) “Bimolecular fluorescence complementation (BiFC): A 5-year update and future perspectives,” BioTechniques, 53(5), pp. 285–298. doi:10.2144/000113943.

Körber, N. (2022) “MIA: An Open Source Standalone Deep Learning Application for Microscopic Image Analysis,” bioRxiv, p. 2022.01.14.476308. doi:10.1101/2022.01.14.476308.

Krull, A., Buchholz, T.-O. and Jug, F. (2019) “Noise2Void - Learning Denoising from Single Noisy Images,” 2019 IEEE/CVF Conference on Computer Vision and Pattern Recognition (CVPR), 00, pp. 2124–2132. doi:10.1109/cvpr.2019.00223.

Laine, R.F. et al. (2022) “High-fidelity 3D live-cell nanoscopy through data-driven enhanced super-resolution radial fluctuation,” bioRxiv, p. 2022.04.07.487490. doi:10.1101/2022.04.07.487490.

LeCun, K.G. and Y. (2021) “Learning Fast Approximations of Sparse Coding,” Proceedings of the 27th International Conference on International Conference on Machine Learning (ICML’10) [Preprint].

Lelek, M. et al. (2021) “Single-molecule localization microscopy,” Nature Reviews Methods Primers, 1(1), p. 39. doi:10.1038/s43586-021-00038-x.

Leonardis, A. and Bischof, H. (2000) “Robust Recognition Using Eigenimages,” Computer Vision and Image Understanding, 78(1), pp. 99–118. doi:10.1006/cviu.1999.0830.

Li, H. et al. (2020) “Live-SIMBA: an ImageJ plug-in for the universal and accelerated single molecule-guided Bayesian localization super resolution microscopy (SIMBA) method,” Biomedical Optics Express, 11(10), p. 5842. doi:10.1364/boe.404820.

Li, Y. et al. (2018) “DLBI: deep learning guided Bayesian inference for structure reconstruction of super-resolution fluorescence microscopy,” Bioinformatics, 34(13), pp. i284–i294. doi:10.1093/bioinformatics/bty241.

Lichtman, J.W. and Conchello, J.-A. (2005) “Fluorescence microscopy,” Nature Methods, 2(12), pp. 910–919. doi:10.1038/nmeth817.

Lin, Y. et al. (2015) “Quantifying and Optimizing Single-Molecule Switching Nanoscopy at High Speeds,” PLoS ONE, 10(5), p. e0128135. doi:10.1371/journal.pone.0128135.

Marsh, R.J. et al. (2021) “Sub-diffraction error mapping for localisation microscopy images,” Nature Communications, 12(1), p. 5611. doi:10.1038/s41467-021-25812-z.

Mendel, J.M. (1991) “Tutorial on higher-order statistics (spectra) in signal processing and system theory: theoretical results and some applications,” Proceedings of the IEEE, 79(3), pp. 278–305. doi:10.1109/5.75086.

Minoshima, M. and Kikuchi, K. (2017) “Photostable and photoswitching fluorescent dyes for super-resolution imaging,” JBIC Journal of Biological Inorganic Chemistry, 22(5), pp. 639–652. doi:10.1007/s00775-016-1435-y.

Moeyaert, B., Vandenberg, W. and Dedecker, P. (2020) “SOFIevaluator: a strategy for the quantitative quality assessment of SOFI data,” Biomedical Optics Express, 11(2), p. 636. doi:10.1364/boe.382278.

Monwar, Md.M., Rezaei, S. and Prkachin, K. (2007) “Eigenimage Based Pain Expression Recognition.” IAENG International Journal of Applied Mathematics.

Nehme, E. et al. (2018) “Deep-STORM: super-resolution single-molecule microscopy by deep learning,” Optica, 5(4), p. 458. doi:10.1364/optica.5.000458.

Opstad, I.S. et al. (2020) “Fluorescence fluctuations-based super-resolution microscopy techniques: an experimental comparative study,” arXiv [Preprint].

Ouyang, W. et al. (2018) “Deep learning massively accelerates super-resolution localization microscopy,” Nature Biotechnology, 36(5), pp. 460–468. doi:10.1038/nbt.4106.

Pawlowska, M. et al. (2022) “Embracing the uncertainty: the evolution of SOFI into a diverse family of fluctuation-based super-resolution microscopy methods,” Journal of Physics: Photonics, 4(1), p. 012002. doi:10.1088/2515-7647/ac3838.

Rao, S., Martins, A. de M. and Príncipe, J.C. (2009) “Mean shift: An information theoretic perspective,” Pattern Recognition Letters, 30(3), pp. 222–230. doi:10.1016/j.patrec.2008.09.011.

Rayleigh, Lord (1903) “XV. On the theory of optical images, with special reference to the microscope,” Journal of the Royal Microscopical Society, 23(4), pp. 447–473. doi:10.1111/j.1365-2818.1903.tb04830.x.

Ronneberger, O., Fischer, P. and Brox, T. (2015) “Medical Image Computing and Computer-Assisted Intervention – MICCAI 2015, 18th International Conference, Munich, Germany, October 5-9, 2015, Proceedings, Part III,” Lecture Notes in Computer Science, pp. 234–241. doi:10.1007/978-3-319-24574-4_28.

Rosten, E., Jones, G.E. and Cox, S. (2013) “ImageJ plug-in for Bayesian analysis of blinking and bleaching.,” Nature methods, 10(2), pp. 97–8. doi:10.1038/nmeth.2342.

Rozell, C.J. et al. (2008) “Sparse Coding via Thresholding and Local Competition in Neural Circuits,” Neural Computation, 20(10), pp. 2526–2563. doi:10.1162/neco.2008.03-07-486.

Rust, M.J., Bates, M. and Zhuang, X. (2006) “Sub-diffraction-limit imaging by stochastic optical reconstruction microscopy (STORM),” Nature Methods, 3(10), pp. 793–796. doi:10.1038/nmeth929.

Sage, D. et al. (2019) “Super-resolution fight club: assessment of 2D and 3D single-molecule localization microscopy software,” Nature Methods, 16(5), pp. 387–395. doi:10.1038/s41592-019-0364-4.

Schermelleh, L. et al. (2019) “Super-resolution microscopy demystified,” Nature Cell Biology, 21(1), pp. 72–84. doi:10.1038/s41556-018-0251-8.

Schnitzbauer, J. et al. (2017) “Super-resolution microscopy with DNA-PAINT,” Nature Protocols, 12(6), pp. 1198–1228. doi:10.1038/nprot.2017.024.

Sibarita, J.-B. (2005) “Deconvolution Microscopy,” Advances in Biochemical Engineering/Biotechnology, pp. 201–243. doi:10.1007/b102215.

Solomon, O. et al. (2018) “Sparsity-based super-resolution microscopy from correlation information,” Optics Express, 26(14), p. 18238. doi:10.1364/oe.26.018238.

Stewart, G.W. (1993) “On the Early History of the Singular Value Decomposition,” SIAM Review, 35(4), pp. 551–566. doi:10.1137/1035134.

Stubb, A. et al. (2020) “Fluctuation-Based Super-Resolution Traction Force Microscopy,” Nano Letters, 20(4), pp. 2230–2245. doi:10.1021/acs.nanolett.9b04083.

Suárez, Y.G. et al. (2019) “Nanoscale organization of rotavirus replication machineries,” eLife, 8, p. e42906. doi:10.7554/elife.42906.

Turk, M. and Pentland, A. (1991) “Eigenfaces for Recognition,” Journal of Cognitive Neuroscience [Preprint]. doi:https://doi.org/10.1162/jocn.1991.3.1.71.

Vangindertael, J. et al. (2018) “An introduction to optical super-resolution microscopy for the adventurous biologist,” Methods and Applications in Fluorescence, 6(2), p. 022003. doi:10.1088/2050-6120/aaae0c.

Wang, B. et al. (2020) “Multicomposite super-resolution microscopy: Enhanced Airyscan resolution with radial fluctuation and sample expansions,” Journal of Biophotonics, 13(5), p. e2419. doi:10.1002/jbio.201960211.

Wegel, E. et al. (2016) “Imaging cellular structures in super-resolution with SIM, STED and Localisation Microscopy: A practical comparison,” Scientific Reports, 6(1), p. 27290. doi:10.1038/srep27290.

Xu, F. et al. (2015) “Bayesian localization microscopy based on intensity distribution of fluorophores,” Protein & Cell, 6(3), pp. 211–220. doi:10.1007/s13238-015-0133-9.

Xu, F. et al. (2017) “Live cell single molecule-guided Bayesian localization super resolution microscopy,” Cell Research, 27(5), pp. 713–716. doi:10.1038/cr.2016.160.

Yahiatene, I. et al. (2015) “Entropy-Based Super-Resolution Imaging (ESI): From Disorder to Fine Detail,” ACS Photonics, 2(8), pp. 1049–1056. doi:10.1021/acsphotonics.5b00307.

Yao, L. et al. (2021) “Dynamic Structure of Yeast Septin by Fast Fluctuation-Enhanced Structured Illumination Microscopy,” Microorganisms, 9(11), p. 2255. doi:10.3390/microorganisms9112255.

Yi, X. et al. (2019) “Moments reconstruction and local dynamic range compression of high order superresolution optical fluctuation imaging,” Biomedical Optics Express, 10(5), p. 2430. doi:10.1364/boe.10.002430.

